# Functional gene delivery to and across brain vasculature of systemic AAVs with endothelial-specific tropism in rodents and broad tropism in primates

**DOI:** 10.1101/2023.01.12.523844

**Authors:** Xinhong Chen, Damien A. Wolfe, Dhanesh Sivadasan Bindu, Mengying Zhang, Naz Taskin, David Goertsen, Timothy F. Shay, Erin Sullivan, Sheng-Fu Huang, Sripriya Ravindra Kumar, Cynthia M. Arokiaraj, Viktor Plattner, Lillian J. Campos, John Mich, Deja Monet, Victoria Ngo, Xiaozhe Ding, Victoria Omstead, Natalie Weed, Yeme Bishaw, Bryan Gore, Ed S Lein, Athena Akrami, Cory Miller, Boaz P. Levi, Annika Keller, Jonathan T. Ting, Andrew S. Fox, Cagla Eroglu, Viviana Gradinaru

## Abstract

Delivering genes to and across the brain vasculature efficiently and specifically across species remains a critical challenge for addressing neurological diseases. We have evolved adeno-associated virus (AAV9) capsids into vectors that transduce brain endothelial cells specifically and efficiently following systemic administration in wild-type mice with diverse genetic backgrounds and rats. These AAVs also exhibit superior transduction of the CNS across non-human primates (marmosets and rhesus macaques), and *ex vivo* human brain slices although the endothelial tropism is not conserved across species. The capsid modifications translate from AAV9 to other serotypes such as AAV1 and AAV-DJ, enabling serotype switching for sequential AAV administration in mice. We demonstrate that the endothelial specific mouse capsids can be used to genetically engineer the blood-brain barrier by transforming the mouse brain vasculature into a functional biofactory. Vasculature-secreted Hevin (a synaptogenic protein) rescued synaptic deficits in a mouse model.

## MAIN

Malfunction of cell types comprising vasculature within the brain, including endothelial cells, can facilitate the progression of neurological disorders (Yu, Ji, and Shao 2020; Xiao et al. 2020; Profaci et al. 2020; Yang et al. 2021). However, the study of vasculature is hampered by limited options for versatile, cell-type-specific transgene delivery. Adeno-associated virus (AAV) vectors offer promise for gene delivery to the brain, but are commonly administered via intracranial injections, resulting in tissue damage and limited and uneven spatial coverage (Aschauer, Kreuz, and Rumpel 2013; Watakabe et al. 2015). Systemic AAV delivery provides a non-invasive, brain-wide alternative (Deverman et al. 2018; Hudry and Vandenberghe 2019; Chen et al. 2022; Challis et al. 2022), but specifically targeting cell types of interest remains a challenge.

Having previously engineered vectors that efficiently cross the blood-brain barrier (BBB) with broad tropism in rodents (e.g. AAV-PHP.eB) (Deverman et al. 2016; Chan et al. 2017; Ravindra Kumar et al. 2020; Goertsen et al. 2021), we recently turned our focus to identifying vectors with cell-type biased tropism. From the PHP.B sequence family, we identified PHP.V1, which, when intravenously delivered, has enhanced potency for endothelial cells although it also transduces astrocytes and neurons (Ravindra Kumar et al. 2020). While an improvement, PHP.V1 still requires cell type-specific promoters whose large size limits the choice of transgenes and only work in certain mice strains. In addition, capsid entry into other cell types may induce an immune response, creating a confounding effect. Similar problems also apply to the previously-reported BR1 endothelial variant, BR1 transduces a large number of neurons and astrocytes other than endothelial cells (Körbelin et al. 2016; Sundaram et al. 2021). With the increasing numbers of engineered AAVs reported, there are also increasing examples showing their potential distinct tropism across species (Chen et al. 2022; Challis et al. 2022), emphasizing the necessity for the testing of engineered variants in different species especially non-human primates.

Another challenge remaining for AAV gene delivery is successful re-administration, which may be needed to maximize therapeutic effect, particularly given the loss of transgene expression over time observed with AAV gene delivery (Greig et al. 2022). While neutralizing antibodies induced by initial AAV administration can prevent sequential administration of the same AAV (Hamilton and Wright 2021), switching to another AAV serotype with similar or complementary features is a potential solution (Bočkor et al. 2017; Colella, Ronzitti, and Mingozzi 2018) that remains underexplored.

An AAV with potent and specific tropism for brain vasculature would enable new strategies for gene therapy. As the leading platform for *in vivo* delivery of gene therapies, AAV vectors have been used to deliver diverse therapeutic genes to treat a broad spectrum of disorders (Lykken et al. 2018; Wang, Tai, and Gao 2019), including those resulting from loss of either cell-autonomous factors or factors which act on neighboring cells regardless of genotype. For disorders caused by the loss of function of a single non-cell-autonomous factor, such as mucopolysaccharidosis (Sawamoto et al. 2018), the factor is typically a secreted protein (Lykken et al. 2018). In these cases, gene therapy could target a healthy cell population, transforming those cells into a ‘biofactory’ for production and secretion of a therapeutic protein that could cross-correct affected cells. Currently, however, therapeutic proteins produced from peripheral biofactories, most commonly the liver, enter the CNS with low efficiency and fall short of rescuing phenotypes in CNS disorders (Ponder and Haskins 2007; Zapolnik and Pyrkosz 2021). Given its broad distribution and close proximity to other cell types within the CNS, AAV-transformed brain vasculature may serve as a better biofactory for the CNS.

Here, through a combination of directed evolution and semi-rational engineering, we identified a family of novel vectors, including AAV-X1 and AAV-X1.1, which target brain endothelial cells specifically and efficiently following systemic delivery in mice with a ubiquitous promoter. We characterized these novel AAVs across rodent models (genetically diverse mouse strains and rats), non-human primates (marmosets and rhesus macaques), and *ex vivo* human brain slices, demonstrating brain endothelial-specific cell targeting in rodent and broad CNS targeting in primates. To illustrate the utility of AAV-X1 for CNS delivery of neuroactive proteins, we transformed mouse brain endothelial cells into a biofactory for producing the synaptogenic protein Sparc-like protein 1 (Sparcl1)/Hevin. Hevin/Sparcl1 is an astrocyte-secreted protein that controls formation of vesicular glutamate transporter 2 (VGluT2)-containing synapses such as thalamocortical synapses (Kucukdereli et al. 2011; Risher et al. 2014; Singh et al. 2016). AAV-X1-mediated ectopic expression of Sparcl1/Hevin in brain endothelial cells was sufficient to rescue the thalamocortical synaptic loss phenotype of Sparcl1/Hevin knockout mice. We also demonstrated the transferability of AAV-X1’s properties from the AAV9 serotype to AAV1, enabling repeated AAV administration to increase CNS transduction. In general, we provide novel engineered systemic AAVs for brain endothelial-specific cell targeting in rodent and broad CNS targeting from rodent to primate with potential for re-administration.

### A novel AAV capsid specifically targets brain endothelial cells in mice following systemic administration

As a starting point for engineering, we chose AAV9, which targets the CNS with low efficiency following systemic delivery. We inserted randomized 7-mer peptides between positions 588 and 589 of AAV9 and injected the resulting virus library intravenously into transgenic mice expressing Cre from either the Tek (targeting endothelial cells) or Synapsin I (Syn, targeting neuronal cells) promoter. After 2 rounds of M-CREATE selection (Ravindra Kumar et al. 2020), we identified a variant, named AAV-X1, which is among the most enriched in Tek-Cre mice, indicating potential endothelial tropism in mice (Fig. 1A). To characterize the transduction profile of AAV-X1 *in vivo* we packaged it with a single-stranded (ss) AAV genome carrying a strong ubiquitous promoter, CAG, driving the expression of an eGFP reporter. Three weeks after IV administration to C57/BL/6J mice, we observed specific and efficient targeting of the vasculature across brain regions (Fig. 1B). To compare the vascular targeting of AAV-X1 to that of previously engineered capsids, we repeated the experiment along with AAV9, AAV-PHP.V1, and AAV-BR1 (Körbelin et al. 2016; Ravindra Kumar et al. 2020). After 3 weeks of expression, AAV-X1 exhibited higher efficiency and higher specificity in targeting brain endothelial cells compared to all 3 controls (Fig. 1C). At a dose of 3 × 10^11^ viral genomes (vg) per mouse, AAV-X1 transduced ∼65-70% of GLUT1+ (endothelial markers) cells across the cortex, hippocampus and thalamus. In comparison, AAV-PHP.V1 and AAV-BR1 transduced ∼40% of GLUT1+ cells, while AAV9 transduced ∼1% of the endothelial cells. AAV-X1 also exhibited superior specificity towards vasculature; ∼95% of the cells transduced by AAV-X1 were GLUT1+. In contrast, AAV-BR1 and AAV-PHP.V1 transduced GLUT1+ cells with much less specificity, with only 60% and 40% of transduced cells being GLUT1+, respectively. AAV-BR1 also targeted neurons, while AAV-PHP.V1 also targeted neurons and astrocytes (Fig. 1C). Further staining with CD31 or Podocalyxin (endothelial cells), Calponin 1 (smooth muscle cells), CD206 (perivascular macrophages), GFAP (astrocytes) and CD13 (pericytes) confirmed the high efficiency and specificity of AAV-X1 in targeting brain endothelial cells (Supplementary Fig. 1A-D). Interestingly, the endothelial cells in the choroid plexus (CP) were not transduced by AAV-X1 (Supplementary Fig. 1C). In summary, the novel vector AAV-X1 not only exhibits significant improvement in targeting the CNS compared to its parent AAV9 but also shows high specificity towards brain endothelial cells.

**Figure 1:**
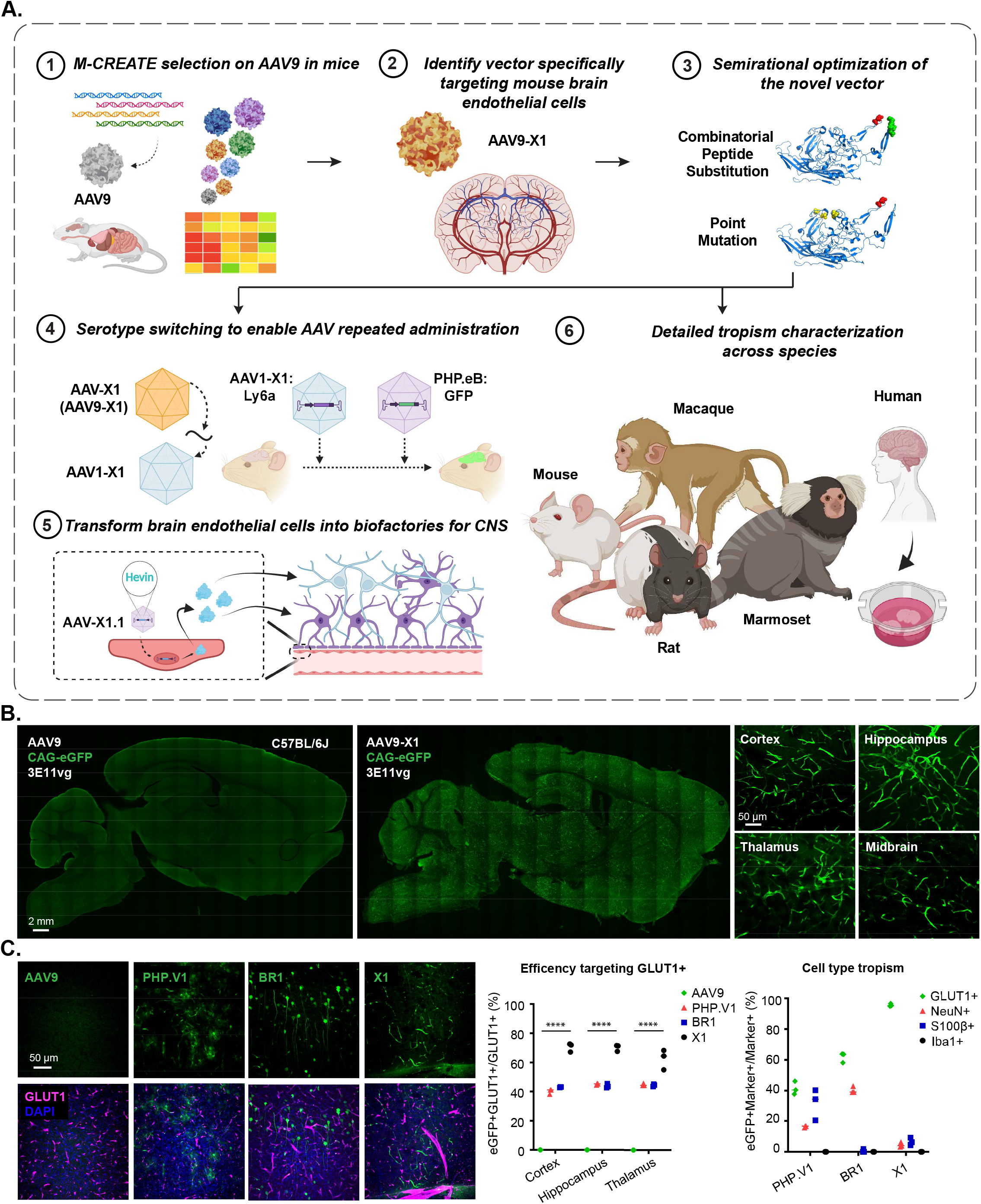
Engineered AAVs can specifically target the brain endothelial cells in mouse following systemic delivery. **A**. An overview of the engineering and characterization of the novel capsids. (1), (2): Evolution of AAV9 using Multiplexed-CREATE and identification of a novel vector, AAV-X1, that transduces brain endothelial cells specifically and efficiently following systemic administration in mice. (3): Combinatorial peptide substitution and point mutation to further refine the novel vector’s tropism, yielding improved vectors. (4): Transferring the X1 peptide to the AAV1 backbone to enable serotype switching for sequential AAV administration. (5): Utilizing AAV-X1 to transform the brain endothelial cells into a biofactory, producing Hevin for the CNS. (6): Validation of the novel AAVs across rodent models (genetically diverse mice strains and rats), NHPs (marmosets and macaques), and *ex vivo* human brain slices. **B**. (Left) Representative images of AAV (AAV9, AAV-X1) vector-mediated expression of eGFP (green) in the brain (scale bar: 2 mm). (Right) Zoomed-in images of AAV-X1-mediated expression of eGFP (green) across brain regions including the cortex, hippocampus, thalamus, and midbrain (scale bar: 50 µm). (C57BL/6J, n=3 per group, 3E11 vg IV dose per mouse, 3 weeks of expression). **C**. (Left) Representative images of AAV (AAV9, PHP.V1, BR1, AAV-X1) vector-mediated expression of eGFP (green) in the cortex. Tissues were co-stained with GLUT1 (magenta) (scale bar: 50 µm). (Middle) Percentage of AAV-mediated eGFP-expressing cells that overlap with the GLUT1+ marker across brain regions, representing the efficiency of the vectors’ targeting of GLUT1+ cells. A two-way ANOVA and Tukey’s multiple comparisons tests with adjusted P values are reported (****P<0.0001 for AAV9 versus X11 in cortex, hippocampus, and thalamus). Each data point shows the mean ± s.e.m of 3 slices per mouse. (Right) Percentage of GLUT1+ markers in AAV-mediated eGFP-expressing cells across brain regions, representing the specificity of the vectors’ targeting of GLUT1+ cells. (C57BL/6J, n=3 per group, 3E11 vg IV dose per mouse, 3 weeks of expression).

### Semi-rational refinement of X1’s peripheral tropism by further engineering cargo and capsid

While the novel variant X1 exhibited significant improvement in transducing the CNS endothelium compared to AAV9, it maintained a similar level of liver transduction. Therefore, we next explored strategies to further refine the vector’s tropism, either by incorporating specific cargo elements or by further engineering the capsid (Fig. 2A).

**Figure 2:**
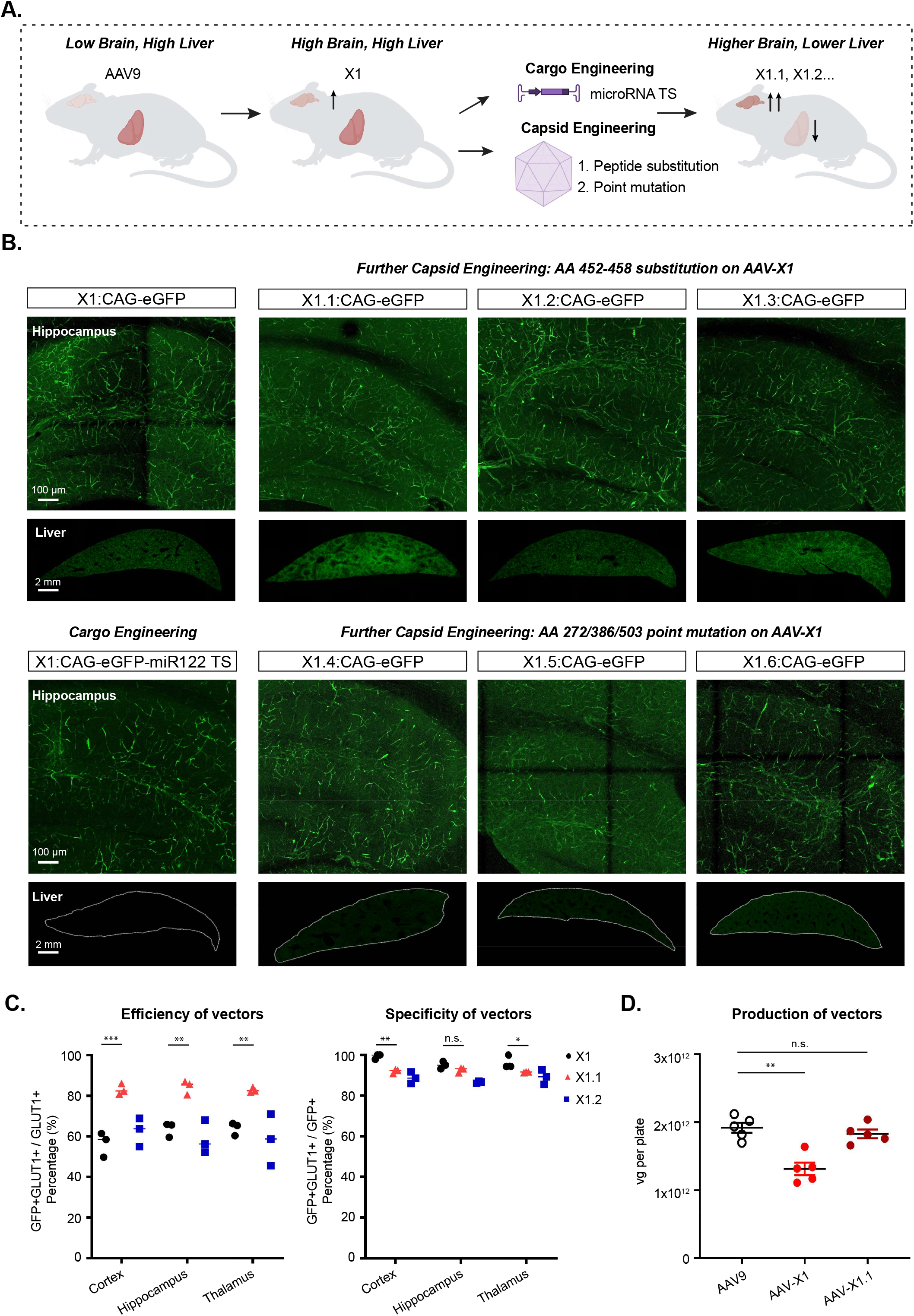
Semi-rational refinement of X1 capsid’s tropism by further cargo and capsid engineering. **A**. Illustration demonstrating cargo and capsid engineering to refine X1’s tropism to increase brain targeting and lower liver targeting. **B**. Representative images of novel vector-mediated expression of eGFP (green) in hippocampus and liver. Images are matched in fluorescence intensity to the X1: CAG-eGFP image. Brain scale bar: 100 µm. Liver scale bar: 2mm. (n≥4 per group, ∼8 week-old C57BL/6J males, 3×10^11^ vg IV dose per mouse, 3 weeks of expression). (Top left) X1-mediated expression of CAG-eGFP. (Bottom left) X1-mediated eGFP expression with cargo engineering by incorporating MicroRNA-122 target sites (miR-122TS) in the CAG-eGFP genome. (Top right) Further capsid engineering by substitution at AA 452-458 of AAV-X1 yielding X1.1, X1.2 and X1.3. Vector-mediated expression of eGFP is shown. (Bottom right) Further capsid engineering on AAV-X1 by mutating AA272/AA386/AA503 to alanine to yield X1.4, X1.6, and X1.5, respectively. Vector-mediated expression of eGFP is shown. **C**. (Left) Percentage of AAV-mediated eGFP-expressing cells that overlap with GLUT1+ markers across brain regions, representing the efficiency of the vectors’ targeting of GLUT1+ cells. A two-way ANOVA and Tukey’s multiple comparisons tests with adjusted P values are reported (P=0.0002 for X1 versus X1.1 in the cortex, P=0.0022 for X1 versus X1.1 in the hippocampus, P=0.0049 for X1 versus X1.1 in the thalamus; *P ≤ 0.05, **P ≤ 0.01 and ***P ≤ 0.001 are shown, P > 0.05 is not shown). Each data point shows the mean ± s.e.m of 3 slices per mouse. (Right) Percentage of GLUT1+ markers in AAV-mediated eGFP-expressing cells across brain regions, representing the specificity of the vectors’ targeting of GLUT1+ cells. A two-way ANOVA and Tukey’s multiple comparisons tests with adjusted P values are reported (P=0.0013 for X1 versus X1.1 in the cortex, P=0.3854 for X1 versus X1.1 in the hippocampus, P=0.0413 for X1 versus X1.1 in the thalamus; *P ≤ 0.05, **P ≤ 0.01, ***P ≤ 0.001, n.s. P > 0.05). Each data point shows the mean ± s.e.m of 3 slices per mouse. **D**. AAV vector yields from an established laboratory protocol (see Methods). One-way analysis of variance (ANOVA) non-parametric Kruskal-Wallis test (approximate P=0.0014), and follow-up multiple comparisons with uncorrected Dunn’s test are reported (P=0.0099 for AAV9 versus AAV-X1, P>0.9999 for AAV9 versus AAV-X1.1; n≥4 per group, each data point is the mean of 3 technical replicates, mean ± s.e.m is plotted). **P ≤ 0.01, n.s. P > 0.05.

We first explored whether adding a regulatory element to X1’s cargo could reduce liver expression. When incorporated into the genome, microRNA target sites induce microRNA-mediated post-transcriptional transgene silencing (Anja Geisler and Fechner 2016). MicroRNA-122 (miR-122) is highly specific to the liver (Isakova et al. 2020) and incorporation of its target site has been shown to reduce AAV expression in the liver (A. Geisler et al. 2011). We cloned 3 copies of miR-122’s target site (TS) into a CAG-eGFP genome and packaged it in the X1 vector. Three weeks after IV delivery of X1: CAG-eGFP-miR122TS, we observed a large reduction in liver expression, indicating that future applications of X1 could incorporate miR-122 TS for liver detargeting (Fig. 2B, bottom left panel).

While cargo engineering reduced transgene expression in the liver, the AAV capsid still entered the liver and might trigger an immune response. Therefore, we next explored whether further engineering of the X1 capsid could achieve similar liver detargeting, and potentially further boost CNS transduction. Structural information (DiMattia et al. 2012) and previous engineering efforts (Goertsen et al. 2021) suggest that the 455 loop of AAV9 potentially interacts with the 588 loop and affects receptor binding. A 7-mer substitution of 452-458 of PHP.eB was found to further boost that capsid’s CNS targeting ability and enable potent transduction of the mouse and marmoset CNS while detargeting the AAV from the liver (Goertsen et al. 2021). We therefore took three 7-mer peptides identified from that PHP.eB 455 loop selection and substituted them into X1, creating X1.1, X1.2, and X1.3. Compared to X1, X1.1 showed further improvement in targeting the CNS, transducing ∼82-85% of GLUT1+ cells across brain regions (Fig. 2B-C). X1.1 also maintained high specificity in targeting the brain vasculature; ∼90-92% of transduced cells were GLUT1+ (Fig. 2C, right). X1.1 showed a similar expression pattern as X1 in peripheral organ and both weren’t vasculature-tropic in the peripheral organs (Supplementary Fig. 2A-B). To fully capture X1.1’s performance across organs, we measured its vector genome and RNA transcript following systemic administration in mice. We observed X1.1’s increased presence in CNS and X1.1’s reduced targeting of most peripheral organs compared to AAV9 (Supplementary Fig. 2C-F). Some application in mice would require a higher dose of AAVs, we thus injected X1.1 with higher doses and verified that it maintained its high specificity against brain endothelial cells (Supplementary Fig. 2G-H). X1.1 was capable of packaging viral genomes with similar efficiency to AAV9 (Fig. 2D). Given X1.1’s high transduction efficiency of brain endothelial cells, we performed a dye perfusion experiment to evaluate whether AAV transduction may compromise BBB integrity. We did not detect increased leakage in the BBB compared to uninjected mice (Supplementary Fig. 3A).

Having seen that X1 can accommodate additional capsid changes, we next investigated whether we could optimize the vector’s tropism by mutating sites known to mediate receptor interactions and affect tropism. N272 and W503 have been shown to be required for AAV9’s binding to galactose (Pulicherla et al. 2011; Shen et al. 2012; Bell et al. 2012; Adachi et al. 2014), and mutation of W503 has been shown to reduce the liver transduction of AAV9 (Pulicherla et al. 2011). We therefore decided to mutate N272 or W503 of X1 to alanine, yielding X1.4 and X1.5, respectively. S386 of AAV9 and Q386 of AAV1 have also been shown to mediate receptor interaction (Adachi et al. 2014; Zhang et al. 2019), so we mutated S386 of X1 to alanine, yielding X1.6. Following IV delivery of these vectors in mice, we observed that X1.4 and X1.5 showed reduced liver transduction while maintaining their brain-endothelial tropism (Fig. 2B, bottom).

To explore the novel vectors’ potential for application in different physiological contexts, we tested the X1.1 vector in young (2.5 months old) versus aged mice (2.5 years old) and did not observe an obvious difference in CNS transduction after 3 weeks’ expression. We also did not observe an obvious sex difference in its CNS tropism (Supplementary Fig. 3B).

### X1 vectors transduce brain endothelial cells across diverse mouse strains and rats in a Ly6a-independent manner

The previously-engineered AAV-PHP.eB has been shown to rely on improved binding to Ly6a for its increased CNS tropism (Hordeaux et al. 2019; Huang et al. 2019; Batista et al. 2020), and polymorphism of Ly6a across mouse strains contributes to its strain-specific phenotype (Hordeaux et al. 2019; Huang et al. 2019). We therefore explored whether the improved CNS targeting of X1 and its further-engineered versions is also dependent on Ly6a binding. We observed that transient overexpression of Ly6a in HEK cells boosted PHP.eB transduction but had no significant effect on X1 transduction (Supplementary Fig. 3C). Surface Plasmon Resonance (SPR) experiments designed to amplify even weak interactions with Ly6a showed that while both PHP.eB and PHP.V1 exhibited strong binding to Ly6a, explaining the strain-specific phenotype of PHP.V1 (Fig. 3A) (Ravindra Kumar et al. 2020), neither X1 nor X1.1 bound Ly6a (Fig. 3A). These results suggest that X1 and its derivatives utilize a novel mechanism of cell targeting.

**Figure 3:**
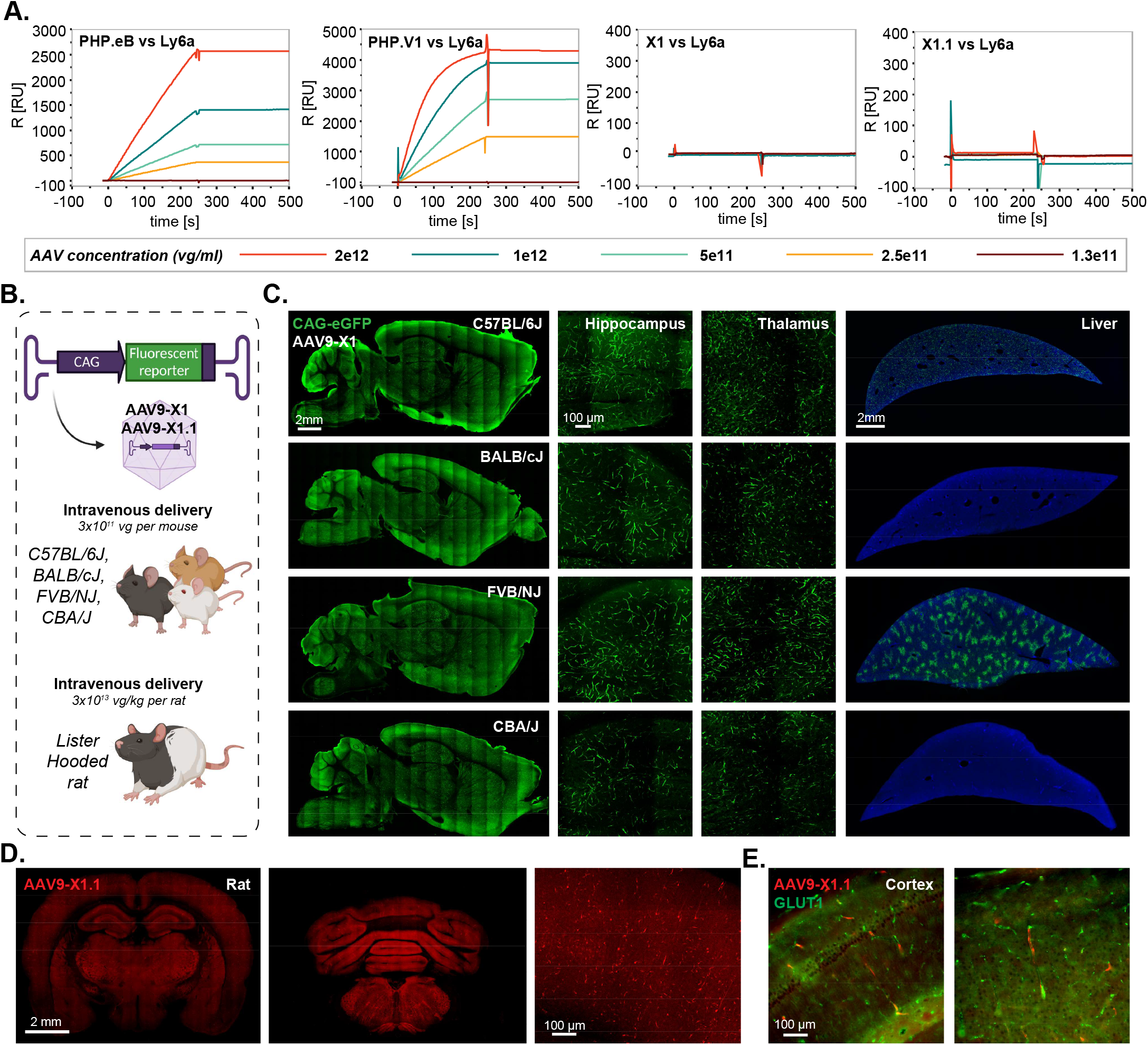
AAV9-X1 and AAV9-X1.1 efficiently transduce brain endothelial cells across diverse mice strains and rats. **A**. Surface plasmon resonance (SPR) plots of PHP.eB, PHP.V1, AAV-X1, and AAV-X1.1 binding to surface-immobilized Ly6a-Fc protein captured on a protein A chip. Colored lines show binding response for each vector across a range of vector concentrations. **B**. Illustration demonstrating the IV administration of AAV-X1 capsid packaged with ssAAV:CAG-eGFP genome in genetically diverse mice strains (∼8 week-old young C57BL/6J, BALB/cJ, FVB/NJ and CBA/J, 3E11 vg per mouse) and IV administration of AAV-X1.1 capsid packaged with ssAAV:CAG-tdTomato genome in Lister Hooded rats (adult, 3E13 vg per rat). **C**. Representative brain and liver images of AAV-X1-mediated eGFP expression (green) in C57BL/6J, BALB/cJ, FVB/NJ and CBA/J mice with zoomed-in images of hippocampus and thalamus. Sagittal brain section scale bar: 2mm. Hippocampus and thalamus scale bar: 100 µm. Liver scale bar: 2mm. **D**. Representative images of forebrain and hindbrain of AAV-X1.1-mediated tdTomato expression (red) in Lister Hooded rat. Scale bar: 2 mm, zoom-in image scale bar: 100 µm. **E**. Representative images of cortex of AAV-X1.1-mediated tdTomato expression (red) in Lister Hooded rat, tissue were co-stained with GLUT1 (green).

The Ly6a-independence of the X1 vectors prompted us to test their translatability across mice strains. Systemic delivery of X1 capsid packaged with ssAAV:CAG-eGFP in BALB/cJ, CBA/J, and FVB/NJ mouse strains resulted in efficient transduction of endothelial cells across brain regions (Fig. 3B,C). Interestingly, though, we observed different expression patterns in the liver across the strains, with minimal liver transduction in BALB/c and CBA/J strains (Fig. 3C).

We next investigated the new variants’ transduction profile in rats. Following intravenous delivery of X1.1 capsid packaging ssAAV:CAG-tdTomato in adult Lister Hooded rats, we observed robust expression in the brain (Fig. 3D). X1.1-induced expression overlapped with GLUT1+, indicating that its tropism towards endothelial cells is conserved (Fig. 3E).

To explore tropism when the BBB is not intact, we next tested the vectors in pericyte-deficient mice (Supplementary Fig. 3D) (Armulik et al. 2010; Mäe et al. 2021). We observed increased transduction of other cell types compared to control mice (Supplementary Fig. 3E). Interestingly, astrocytes with end-feet on the brain vasculature were transduced in pericyte-deficient mice but not in control mice (Supplementary Fig. 3F), indicating that the novel vectors’ specificity towards endothelial cells may be dependent on the status of the BBB.

### Serotype switching of X1 from AAV9 into AAV1 enables repeated administration of AAV, increasing permeability of the mouse CNS to AAVs

Repeated administration of AAV is desired for certain applications, however, neutralizing antibodies induced by an initial administration prevent the subsequent delivery of similar AAVs (Bortolussi et al. 2014; Colella, Ronzitti, and Mingozzi 2018). One potential solution is to switch AAV serotypes in sequential administrations, but most of the novel AAVs engineered to target the CNS, including PHP.B and AAV-X1, are based on AAV9. To overcome this potential hurdle and enable repeated administration, we sought to transfer the X1 peptide and recreate our engineered AAV in another natural serotype.

Mixed results have been reported for previous attempts to directly transfer the PHP.B or PHP.eB peptide to natural serotypes, usually AAV1 due to its CNS tropism (Tan et al. 2019; Lau et al. 2019; Martino et al. 2021; Pietersz et al. 2021). We inserted the 7-mer peptide of PHP.B into the AAV1 capsid, creating the hybrid AAV1-PHP.B (Fig. 4B). Intravenous injection of adult mice with AAV1-PHP.B packaging CAG-eGFP showed no obvious improvement in brain transduction compared to AAV1. However, AAV1-X1, created by a similar strategy, showed robust CNS expression in a pattern similar to that of X1, efficiently and specifically targeting GLUT1+ endothelial cells (Fig. 4C). We also tried tested similar strategy and inserted the X1 peptide in AAV-DJ, creating AAV-DJ-X1. Similar to AAV1-X1, AAV-DJ-X1 showed improved CNS targeting compared to AAV-DJ following systemic delivery (Supplementary Fig. 4A). These results indicate that X1’s phenotype can be transferred to some other serotypes by transferring the 7-mer peptide, while it would need to be tested individually for each new serotype.

**Figure 4:**
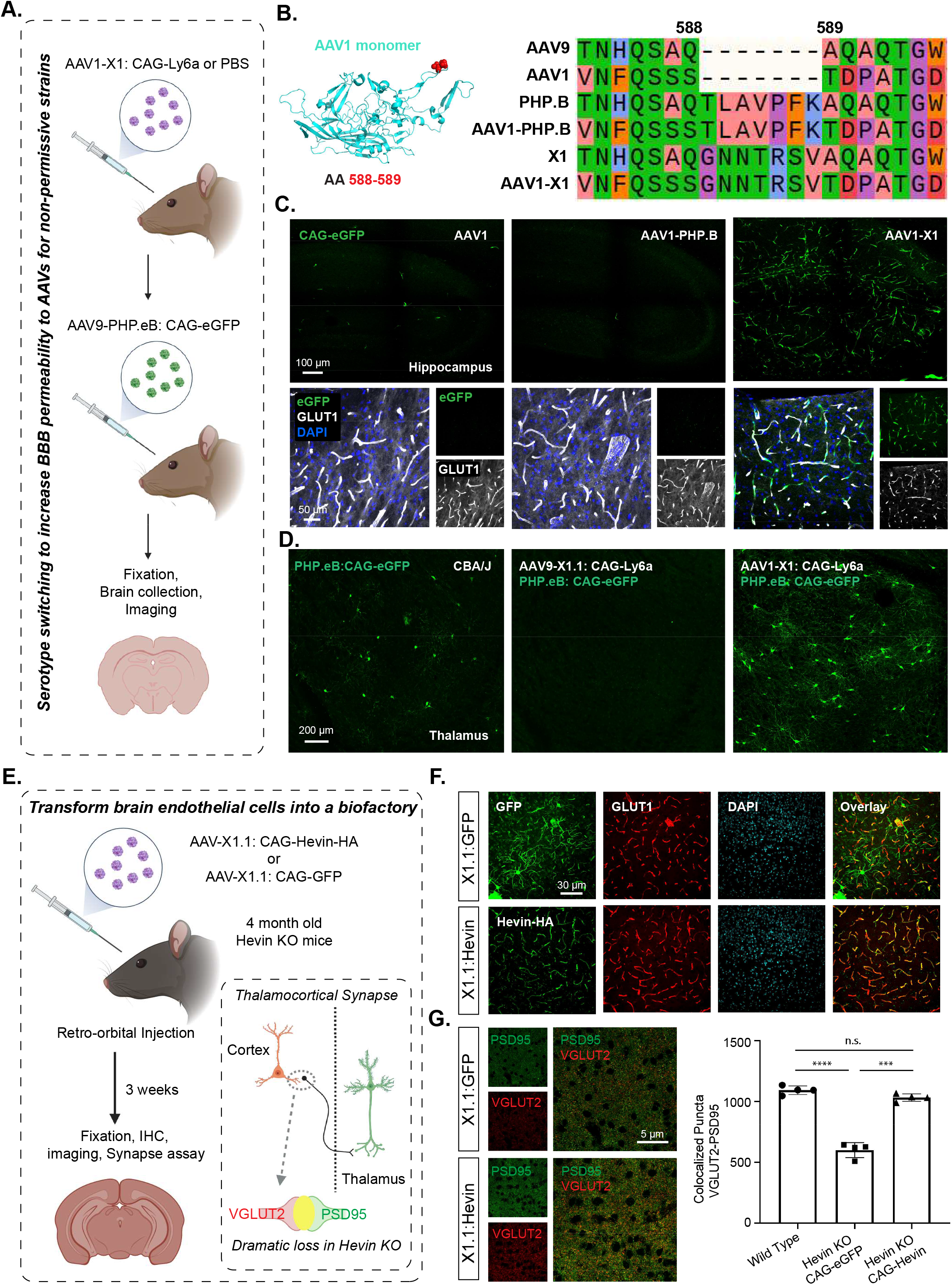
X1 capsids can enable AAV repeated administration with serotype switching and can transform the BBB into a biofactory. **A**. Illustration of utilizing serotype switching to increase BBB permeability for AAVs in non-permissive strains. IV administration of AAV1-X1 capsid packaged with ssAAV: CAG-Ly6a or PBS in CBA/J mice (∼8-week-old young adults, 3×10^11^ vg IV dose/mouse, n=4). After 3 weeks, AAV9-PHP.eB packaged with CAG-eGFP was intravenously administered into CBA/J mice. 3 weeks after the second injection, the brain was collected and imaged. **B**. (Left) Illustration of AAV1 monomer structure with the position of the 7-mer insertion, AA 588/589, highlighted (red). (Right) Sequences of AAV9, AAV1, PHP.B, AAV1-PHP.B, X1 and AAV1-X1 around the insertion site are shown, colored based on physicochemical properties (Zappo). **C**. (Top) Representative images of AAV1, AAV1-PHP.B and AAV-X1-mediated eGFP expression in the hippocampus (scale bar: 100 µm). (Bottom) Zoomed-in images of tissues co-stained with GLUT1 (white) and DAPI (blue). **D**. (Left) Representative image of PHP.eB-mediated eGFP expression in the brain of CAB/J mice (scale bar: 200 μm). (Right) Representative images of the brains of CAB/J mice following sequential administration of either AAV9-X1.1: CAG-Ly6a or AAV1-X1: CAG-Ly6a followed by PHP.eB:CAG-eGFP. **E**. Illustration of transforming brain endothelial cells into a biofactory. IV administration of AAV-X1.1 capsid packaged with ssAAV:CAG-Hevin-HA or ssAAV:CAG-eGFP genome in Hevin-KO mice (∼4-month-old young adults, 1×10^12^ vg IV dose/mouse, n=4). Three weeks post-expression, the mice were anesthetized and perfused and fixation and IHC were performed on the brains. (Bottom right) Illustration of the thalamocortical synapses identified by co-localization of VGLUT2 and PSD95 staining. Thalamocortical synapses are lost in Hevin-KO mice. **F**. (Top) Representative images of AAV-X1.1 vector-mediated expression of eGFP (green) in the brain. The tissues were co-stained with either GLUT1 (red) or DAPI (blue) markers (scale bars: 30 µm). (Bottom) Representative images of AAV-X1.1 vector-mediated expression of Hevin in the brain. The tissues were co-stained with HA (green), GLUT1 (red) and DAPI (blue) markers (scale bar: 30 µm). **G**. (Left) Representative images of a cortical slice stained for PSD95 (green) and VGLUT2 (red) (scale bar: 5 µm). (Right) Quantification of colocalized puncta of VGLUT2 and PSD95 in mice administered X1.1: CAG-eGFP and X1.1: CAG-Hevin-HA. One-way analysis of variance (ANOVA) Brown-Forsythe test (****P<0.0001 for WT versus Hevin KO: CAG-eGFP, ***P=0.0006 for Hevin KO: CAG-eGFP versus Hevin KO: CAG-Hevin, n.s. P=0.1156 for WT versus Hevin KO: CAG-Hevin; n=4 per group, each data point is the mean of 15 slices for one animal, mean ± s.e.m is plotted).

To test AAV1-X1’s potential to enable repeated administration, we utilized the virus for Ly6a supplementation. Polymorphisms in Ly6a in certain mouse strains such as CBA/J and BALB/cJ greatly reduce the CNS permeability of PHP.eB (Hordeaux et al. 2019; Huang et al. 2019; Batista et al. 2020). The brain endothelial cell tropism of X1 and AAV1-X1 prompted us to utilize them to express C57BL/6J-like Ly6a in the BBB of these non-permissive strains, and thereby increase the permeability of those animals’ BBB for AAV9-based PHP.eB when it is subsequently administered. We intravenously delivered AAV1-X1 or X1.1 capsid packaged with Ly6a into adult CBA/J mice. After three weeks of expression, we then injected PHP.eB packaged with eGFP into the same mice (Fig. 4A, D). We observed increased eGFP expression in the brains of CBA/J mice injected with AAV1-X1: CAG-Ly6a but not the mice injected with X1.1: CAG-Ly6a, indicating that AAV1-X1 enables the subsequent administration of PHP.eB and also facilitates the permeability of PHP.eB (Fig. 4D). This result indicates that the serotype switching paradigm enabled by AAV1-X1 could potentially provide a solution for AAV re-administration in mice.

### The X1.1 vector can transform brain endothelial cells into a biofactory for secreted protein delivery to the brain

The broad distribution of vasculature across brain regions creates the opportunity to transform endothelial cells into a biofactory for the broad production of therapeutic agents such as secreted proteins with trophic properties for other cell types within the CNS. For secreted proteins, this would remove the production burden from the target cell, which may already reside in a disease state. Since our novel vectors transduce brain endothelial cells efficiently and specifically, they provide a unique tool for us to explore this concept.

Sparc-like protein 1 (Sparcl1), which is also known as Hevin, is a matricellular secreted protein that is predominantly expressed by astrocytes and a subset of neurons in the CNS. Endothelial cells also express Sparcl1 mRNA; however, protein expression by these cells has not been observed (Lloyd-Burton and Roskams 2012; Mongrédien et al. 2019). Downregulation or missense mutations in SPARCL1/Hevin have been reported in numerous neurological disorders such as autism, schizophrenia and depression (Kähler et al. 2008; Zhurov et al. 2012; De Rubeis et al. 2014). In the developing mouse visual cortex, Hevin is specifically required for the formation and plasticity of thalamocortical connections (Risher et al. 2014; Singh et al. 2016). Hevin knockout mice (Hevin KO) display a dramatic loss of Vesicular Glutamate Transporter 2 positive (VGluT2+) thalamocortical synapses both in the first three postnatal weeks and as adults (Fig. 4E) (Risher et al. 2014).

To determine if ectopic expression of Hevin by CNS endothelial cells can rescue the deficits observed in Hevin KO mice, we generated a viral vector for Hevin expression using our capsid AAV X1.1. Indeed, AAV X1.1 packaging Hevin efficiently transduced brain endothelial cells and drove the expression of Hevin in these cells. To investigate whether Hevin production via endothelial cells would rescue the deficits in thalamocortical synapses observed in Hevin KO mice, we retro-orbitally delivered both Hevin-HA and eGFP driven by the CAG promoter into 4-month-old Hevin KO mice (Fig. 4E). After 3 weeks, we observed robust and specific expression of transgenes in brain endothelial cells (Fig. 4F; Supplementary Figure. 4B-C). Further, a synapse assay using the presynaptic thalamocortical marker VGlut2 and postsynaptic marker PSD95 in Hevin KO mice showed a significantly higher number of thalamocortical synapses in layer IV of the V1 cortex in the treated group (CAG-Hevin) compared to the control (CAG-eGFP) (Fig. 4G). This result indicates that the Hevin produced and secreted from the brain endothelial cells was able to mimic endogenous Hevin function, rescuing the thalamocortical synapse formation that is severely deficient in Hevin KO mice. This system thus illustrates a new potential therapeutic strategy for Hevin-related disorders.

### The X1 vector family efficiently transduces human brain endothelial cells *in vitro*

After validating the novel vectors’ transduction profiles and demonstrating their functional application in rodent species, we next explored translation to higher organisms. We measured the performance of X1 in transducing isolated human brain microvascular endothelial cells (HBMECs). At an MOI of 3 × 10^4^, X1 transduced ∼42% of HBMECs, ∼180 fold higher than its parent AAV9. The further engineered versions X1.1 and X1.2 transduced 44% and 53% of HBMECs, respectively. In contrast, previously-engineered AAV vectors including PHP.eB and BR1 transduced ∼0-2% of HBMECs, showing no obvious improvement in transduction compared to natural serotypes (Fig. 5A). AAV1-X1 and AAV-DJ-X1 also showed improvement in transduction pf HBMECs compared to their parent serotype, respectively (Supplementary Figure. 4D). We next examined the vectors’ performance at a lower MOI of 3 × 10^3^ and again observed an improvement in transduction efficiency for X1.1 and X1.2, which transduced ∼22% and ∼28% of HBMECs (Fig. 5A). In addition to transducing HBMECs, X1 and its engineered derivatives exhibited robust transduction of multiple other human-derived cell lines, including HeLa, U87 and IMR-32 (Supplementary Fig. 4A).

**Figure 5:**
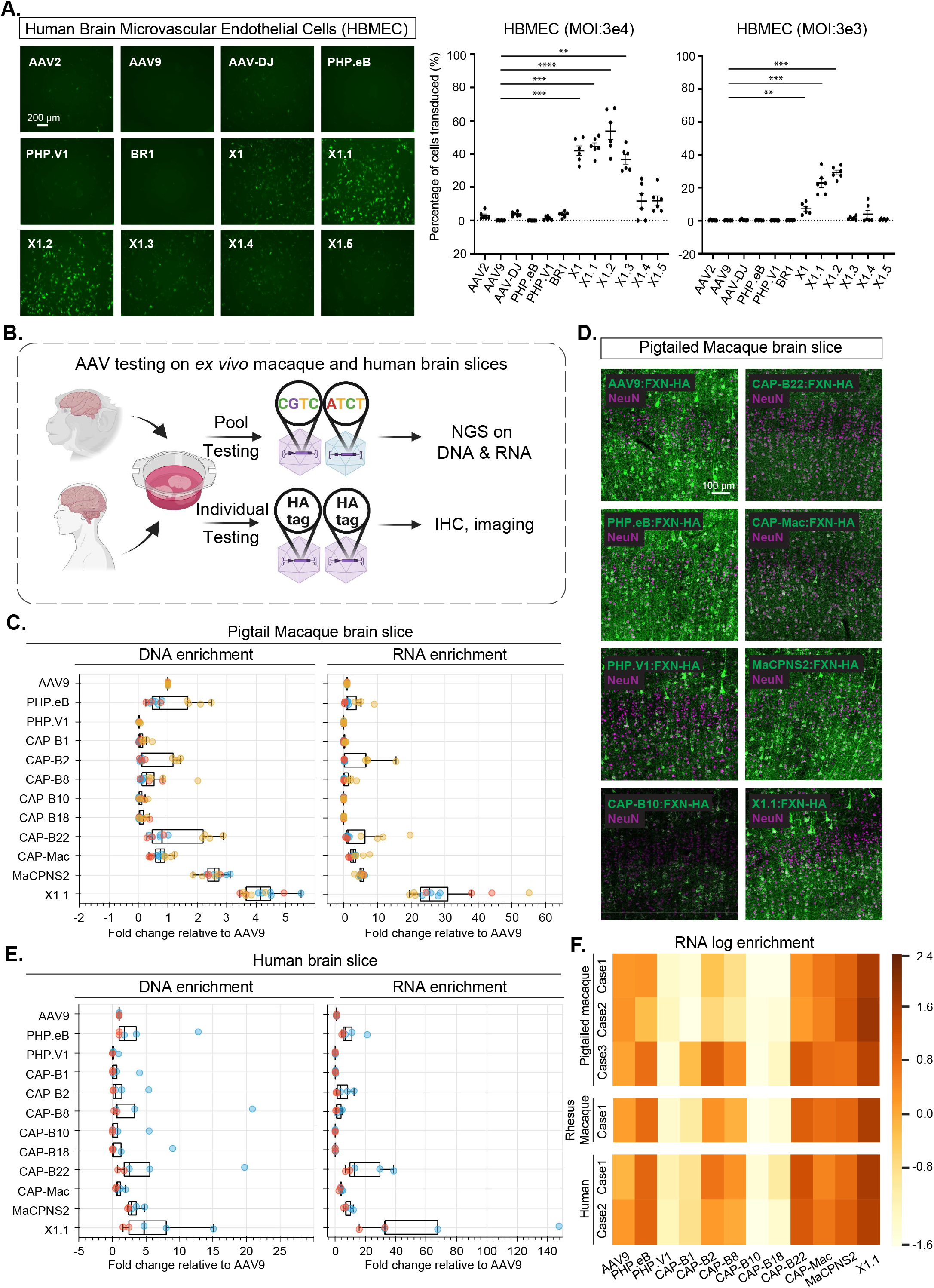
Engineered AAVs can efficiently transduce cultured human brain microvascular endothelial cells, *ex vivo* macaque and human brain slices. **A**. (Left) Representative images of AAV (AAV2, AAV9, AAV-DJ, PHP.eB, PHP.V1, BR1, X1, X1.1, X1.2, X1.3, X1.4, X1.5)-mediated eGFP expression (green) in Human Brain Microvascular Endothelial Cells (HBMECs). (AAVs packaged with ssAAV:CAG-eGFP, n= 6 per condition, 1 day expression). (Right) Percentage of cells transduced by the AAVs. In the condition of MOI:3E4, one-way analysis of variance (ANOVA) non-parametric Kruskal-Wallis test (approximate P<0.0001), and follow-up multiple comparisons with uncorrected Dunn’s test are reported (P=0.0004 for AAV9 versus AAV-X1, P=0.0002 for AAV9 versus AAV-X1.1, P<0.0001 for AAV9 versus AAV-X1.2, P=0.002 for AAV9 versus AAV-X1.3; n=6 per group, each data point is the mean of 3 technical replicates, mean ± s.e.m is plotted). In the condition of MOI:3E3, one-way analysis of variance (ANOVA) non-parametric Kruskal-Wallis test (approximate P<0.0001), and follow-up multiple comparisons with uncorrected Dunn’s test are reported (P=0.0082 for AAV9 versus AAV-X1, P=0.0004 for AAV9 versus AAV-X1.1, P=0.0001 for AAV9 versus AAV-X1.2; n=6 per group, each data point is the mean of 3 technical replicates, mean ± s.e.m is plotted). **P ≤ 0.01, ***P ≤ 0.001, ****P ≤ 0.0001 are shown, P > 0.05 is not shown. **B**. Illustration of AAV testing in *ex vivo* macaque and human brain slices. The brain slices were freshly extracted from southern pig-tailed macaque brain, rhesus macaque brain, and human brain. The slices were cultured at physiological conditions *ex vivo*. In the pool testing pipeline (top), a pool of AAVs packaged with CAG-FXN-HA genome containing a unique barcode was applied to the slice, and DNA extraction and RNA extraction were performed after 7 days. Next-generation sequencing (NGS) was performed to determine the proportion of each barcode (AAV) in DNA and RNA. In the individual testing pipeline (bottom), AAVs packaged with CAG-FXN-HA were individually applied to the slices. Fixation, IHC, and imaging were performed on the slices after 7 days. **C**. DNA and RNA level in southern pig-tailed macaque brain slices for AAVs, with DNA and RNA levels normalized to AAV9. **D**. Representative images of AAV-mediated CAG-FXN-HA expression in *ex vivo* southern pig-tailed macaque brain slices. The tissues were co-stained with antibodies against HA (green) and NeuN (magenta). **E**. DNA and RNA level in human brain slices for AAVs, with DNA and RNA levels normalized to AAV9. **F**. RNA log enrichment of AAVs across pigtailed macaque, rhesus macaque and human brain slices.

### X1.1 efficiently transduces cultured *ex vivo* brain slices from macaque and human

The strong performance of the novel vectors in HBMECs prompted us to investigate their efficacy in a system that may better mimic *in vivo* conditions in non-human primates. We thus turned to an *ex vivo* brain slice culture system (Fig. 5B). The brain slices were freshly extracted from primate brains and cultured ex vivo. The versatile nature of this ex vivo system provides us with a rare opportunity to compare most of the CNS-tropic AAVs engineered in our lab within the same condition in primate tissue. We packaged CAG-Frataxin(FXN)-HA with unique 12bp barcodes in a panel of AAVs including AAV9, X1.1, and several other engineered CNS-tropic AAVs. We then injected them both individually and together as a pool into brain slices freshly extracted from the southern pig-tailed macaque (*Macaca nemestrina*) and cultured in physiological conditions (Fig. 5B). After 7-10 days, we extracted DNA and RNA from the tissue, and calculated the enrichments of the variants from the AAV pool in the tissue using next-generation sequencing (NGS). X1.1 showed an ∼3-fold increase and ∼24-fold increase in DNA and RNA, respectively, compared to AAV9. This outperformed all other vectors, including CAP-B10 and CAP-B22, which have been shown to have increased efficiency in transducing the CNS of marmosets after systemic delivery (Goertsen et al. 2021) (Fig. 5C). The previously-reported MaCPNS2, which has increased targeting of the macaque CNS, also showed an ∼5-fold increase in RNA compared to AAV9 (Chen et al. 2022) (Fig. 5C). We also observed a higher DNA and RNA presence of X1.1 compared to other AAVs in the pool in the rhesus macaque brain (Supplementary Fig. 5A-B). Immunohistochemistry (IHC) staining of the HA tag confirmed the robust transduction by X1.1 in pig-tailed macaque (Fig. 5D; Supplementary Fig. 5C). Interestingly, X1.1-induced protein expression in the *ex vivo* brain slices were mostly seen in neurons (Fig. 5D; Supplementary Fig. 5D). RNA scope experiment revealed that some X1.1-deliverered GFP mRNA transcripts were also observed in GLUT1+ cells while most of them remained in neurons (Supplementary Fig. 5E-F).

We then performed similar testing on human brain slices originating from a biopsy surgery. X1.1 showed an ∼4-fold and ∼32-fold increase in DNA and RNA, respectively, compared to AAV9, again outperforming several previously-engineered CNS-targeting vectors (Fig. 5E). The strong performance of X1.1 across macaque and human *ex vivo* slices (Fig. 5F) suggests great potential for the capsid in future translational applications.

### X1.1 efficiently targets the CNS in rhesus macaque following IV delivery

Having validated the new variants’ transduction profiles in *ex vivo* NHP and human brain models, we decided to investigate their efficacy *in vivo*. We first tested them in the marmoset, a New World monkey. To reduce animal subject numbers, we simultaneously tested two viral capsids (AAV9 and X1.1) packaging different fluorescent reporters (either ssAAV:CAG-eGFP or ssAAV:CAG-tdTomato) in each adult marmoset (Supplementary Fig. 6A). 3 weeks after IV delivery of the viruses, we observed no obvious differences between the CNS transduction of X1.1 and AAV9 (Supplementary Fig. 6B-C). This result emphasizes again that BBB heterogeneity should be taken into account when performing AAV engineering.

We then tested their efficacy in the rhesus macaque, an Old World monkey. We intravenously injected a pool of AAV9 and X1.1 each packaged a unique barcode into a neonate rhesus macaque (male *Macaca mulatta*, within 10 days of birth, 1×10^13^ vg/kg per AAV, n=1 per group) (Supplementary Fig. 7A). After 4 weeks of expression, X1.1 showed an increase in vector genome and RNA transcript across brain regions include cortex, hippocampus and cerebellum, compared to AAV9 (Supplementary Fig. 7B-D). In peripheral tissue such as liver, muscle and enteric system, X1.1 showed a reduction in both DNA and RNA compared to AAV9 (Supplementary Fig. 7B-D). After seeing X1.1 superior performance in targeting CNS while detargeting from peripheral at DNA and RNA levels, we next examined its cell-type targeting profile in the CNS. We packaged ssAAV:CAG-eGFP in AAV9 and ssAAV:CAG-tdTomato in X1.1, then intravenously delivered them as an AAV cocktail to a neonate rhesus macaque (female *Macaca mulatta*, within 10 days of birth, 5×10^13^ vg/kg per macaque, 2.5×10^13^ vg/kg per AAV, n=1 per group) (Fig. 6A). After 4 weeks of expression, we observed robust expression of X1.1 across brain regions including cortex, lingual gyrus (LG), hippocampus, and cerebellum, compared to AAV9 (Fig. 6B-C; Supplementary Video. 1). Further IHC staining revealed that ∼98% of the cells transduced by X1.1 in the cortex were neurons, while a small proportion of the targeted cells were endothelial or glia cells (Fig. 6D-E). Neuron transduction by X1.1 was ∼45-fold higher than AAV9 (Fig. 6F). The difference between the neurotropic profile of X1.1 in neonate macaque and its endothelial cell-tropic profile in rodents both opens up new potential applications and highlights the necessity of profiling AAV vectors across species. These experiments demonstrate that the new capsid AAV-X1.1 can efficiently transduce the CNS in infant Old World monkeys, making it a potentially useful vector for research of neurological disorders.

**Figure 6:**
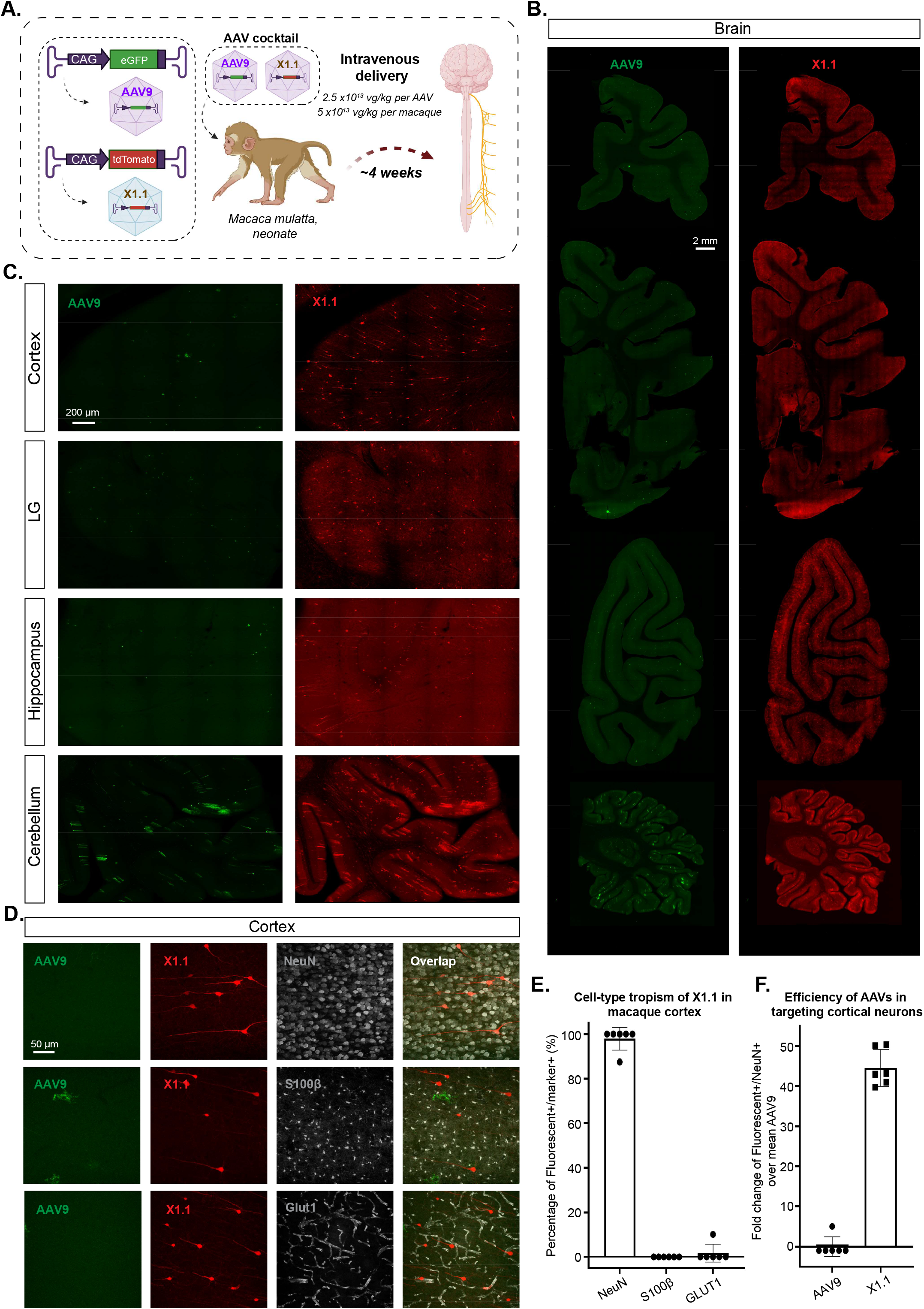
Engineered AAVs can efficiently transduce the central nervous system in rhesus macaque. **A**. Illustration of AAV vector delivery to rhesus macaque to study transduction across the CNS and PNS after 3 weeks of expression. The capsids (AAV9/X1.1) and their corresponding genomes (ssAAV:CAG-eGFP/tdTomato) are shown on the left. Two AAVs packaged with different fluorescent proteins were mixed and intravenously injected at a dose of 5×10^13^ vg/kg per macaque (*Macaca mulatta*, injected within 10 days of birth, female, i.e. 2.5×10^13^ vg/kg per AAV). Representative images of macaque **B**., coronal sections of forebrain, midbrain, hindbrain and cerebellum (scale bar: 2 mm), and **C**., selected brain areas: cortex, lingual gyrus (LG), hippocampus and cerebellum (scale bar: 200 µm). **D**. Brain tissues were co-stained with NeuN (white) or S100β (white) or GLUT1 (white); representative images of the cortex are shown. **E**. Cell-type tropism of X1.1 in macaque brain, shown by percentage of Fluorescent+/Marker+. Each data point is a slice. **F**. Quantification of the fold change of Fluorescent+/NeuN+ over mean AAV9 in the macaque brain. Each data point is a slice.

## DISCUSSION

In this study, we describe a family of novel vectors including AAV-X1 and AAV-X1.1, which specifically and efficiently transduces mouse brain endothelial cells with a ubiquitous promoter following systemic administration. This level of specificity in the mouse CNS is unprecedented among both natural and previously-engineered AAV serotypes. The previously-engineered AAV vector PHP.eB (Chan et al. 2017) has been widely used for targeting most cell types in mice CNS, while the more recently-engineered vector PHP.V1 (Ravindra Kumar et al. 2020) has been shown to have increased potency for, but not selective targeting of, brain endothelial cells. However, the enhanced CNS tropism of both vectors is absent in a subset of mouse strains, including BALB/cJ. Here we show that, similar to PHP.eB, PHP.V1 relies on Ly6a, while AAV-X1 vectors are Ly6a-independent and efficiently target brain endothelial cells across mouse strains. This Ly6a independence suggests that these novel vectors may utilize a novel receptor for CNS targeting and adds to the vectors’ translational promise, as Ly6a is a murine-specific factor.

A major challenge in achieving successful gene therapy is the presence of neutralizing antibodies against AAVs. The neutralizing antibodies induced by an initial AAV delivery have been reported to persist long after the treatment, which could prevent the successful repeated administration needed for maintaining transgene expression (Hamilton and Wright 2021). Serotype switching between administrations could be a potential solution for dealing with neutralizing antibodies against the initially-administered serotype. We successfully transferred the 7-mer X1 peptide from AAV9 to AAV1, yielding AAV1-X1, which transduces brain endothelial cells efficiently following IV delivery. This result shows that the X1 peptide is more modular than the 7-mer peptide of PHP.B (Martino et al. 2021). Administering AAV1-X1 packaging Ly6a, we successfully made CBA/J mice more permissive for PHP.eB, thus both demonstrating the novel vector’s ability to introduce a functional receptor and verifying that serotype switching may be a potential solution for sequential AAV administration (Rivière, Danos, and Douar 2006; Majowicz et al. 2017). The transferability of the AAV9-based X1’s 7-mer peptide to AAV1 and AAV-DJ highlights a general strategy that could be applied to other engineered AAVs, although the previous failure of the transfer of PHP.B’s CNS-tropic phenotype along with its peptide to AAV1 (Tan et al. 2019; Lau et al. 2019; Martino et al. 2021; Pietersz et al. 2021) cautions that the success of this strategy could vary case by case (Supplementary Table 1). Engineered AAVs in diverse serotype backbones could not only help with applications involving repeated administration but also circumvent pre-existing neutralizing antibodies to a certain AAV serotype.

Delivering therapeutic agents, or genetic material encoding therapeutics, across the BBB to treat neurological diseases of the CNS remains a challenging task (Stanimirovic, Sandhu, and Costain 2018; Terstappen et al. 2021). Many enzymes, however, are secreted and can exert cross-correction effects. For these, we could instead deliver the genetic material to brain endothelial cells, transforming these cells into a biofactory to produce and distribute therapeutics to other cell types. The novel endothelial cell-tropic vector we describe here provided us with a unique chance to explore this concept in an animal model. We used the X1.1 vector to induce Sparcl1/Hevin protein production in brain endothelial cells and observed that this rescued the thalamocortical synapse loss phenotype of Hevin KO mice. This proof-of-concept supports the brain endothelial cell biofactory model for production of enzymes, antibodies, or other biological therapeutics, suggesting a novel therapeutic approach for diseases like lysosomal storage disorders.

Previous systemic AAVs for the CNS that were engineered in mice have not always translated to NHPs (Hordeaux et al. 2018; Matsuzaki et al. 2018). Tropism of previously-reported endothelial variants such as BR1 or BI30 (Körbelin et al. 2016; Krolak et al. 2022) remained unexplored in non-human primate. Given the X1 vectors’ independence of Ly6a, we tested the novel vectors in various NHP species. We initially tested X1.1 in marmoset, a New World monkey, but did not observe an obvious difference in targeting the CNS compared to AAV9 following systemic delivery. We then turned to the macaque, an Old World monkey, which is more similar genetically to humans and is widely used in research, including gene therapy (Jennings et al. 2016; Hudry and Vandenberghe 2019; Bey et al. 2020). We first tested the virus on *ex vivo* brain slices from the rhesus macaque (*Macaca mulatta*) and southern pig-tailed macaque (*Macaca nemestrina*). We observed greater increases in DNA, RNA, and protein with X1.1 than other CNS vectors including PHP.eB (Chan et al. 2017), CAP-B10, and CAP-B22 (Goertsen et al. 2021). We then intravenously injected X1.1 in the rhesus macaque and observed a significant improvement in targeting the CNS compared to AAV9. Interestingly, the rodent-endothelial-tropic X1.1 efficiently transduces neurons in the macaque brain. The conservation of enhanced CNS tropism across rodents and NHPs is encouraging for X1.1’s potential in tackling neurological disorders in specific condition. We further showed that our engineered vector X1.1 also has increased efficiency in transducing *ex vivo* adult macaque and human brain slices compared to other previously-engineered CNS vectors. Further experiments would be needed to examine the novel vectors’ translation potential in adult primates.

The exact mechanism behind the distinction in cell-type tropism of X1.1 across models tested remains unknown. The different ages of animals tested, the difference in BBB maturation and the different route of administration would also need to be taken into consideration for this distinction (Supplementary Table 2). In rodents with a healthy BBB, X1 vectors seem to prefer endocytosis to transcytosis at the BBB, either due to their interactions with novel receptors or the vector’s own physiological features. In pericyte-deficient mice where endothelial transcytosis is increased and endothelial cells show occasional hot-spot leakage areas due to altered endothelial cell-cell interaction (Armulik et al. 2010; Mäe et al. 2021), we observed increasing transduction of astrocytes and neurons. In rhesus macaque, X1.1 seems biased towards transcytosis at the BBB and transduces neurons efficiently after crossing the BBB. This distinction in tropism opens up the potential for different applications with X1.1 in endothelial cells and neurons in different species. It also emphasizes the need to test AAV variants across species to capture their complete profile.

In summary, here we describe new systemic AAV tools to expand our understanding of the neurovascular unit across species. The novel vector X1 and its further-engineered family of variants, including X1.1, provide genetic access to brain endothelial cells in mice with unprecedented potency and specificity, and their efficient targeting of the CNS in NHP and human brain slices offers hope of accelerating translational research. The surprising modularity of the X1 variant peptide may allow researchers to apply immunogenically-distinct AAVs at multiple timepoints, and the demonstrated application of the X1 vectors to transform brain endothelial cells into a secretory biofactory validates a novel method to deliver therapeutic agents to the CNS.

## CONTRIBUTIONS

X.C. and V.G. designed the experiments. X.C., D.A.W., J. T.T., M. Z., D.S.B., H.S., S.R.K., T.F.S., E.S., D.G., V.N. performed experiments. D.A.W. assisted with the characterization of the vectors across models. D.S.B. and C.E. designed and conducted the *in vivo* Hevin expression experiments. J.T.T., M.Z., D.G., V.O., N.T., N.W., J.M., Y.B., D.M., B.G., B.P.L and E.S.L assisted with the characterization of variants in *ex vivo* macaque and human brain slices. T.M. and E.S. assisted with the SPR experiment and the Ly6a experiment in HEK293T cells. S.H. and A.K. assisted with the characterization of the vectors in pericyte-deficient mice and vascular cell type specific transduction. S.R.K assisted with the HBMEC experiment and early iteration of the library. C.M.A assisted with tissue processing in macaque. X.Z generated the AAV1 and AAV9 models. V.P. and A.K. assisted with the characterization in rat with the support of the staff at SWC. V.N. and C.T.M. assisted with the characterization of the virus in marmoset with the support of vet staff at UCSD. L.J.C and A.F assisted with the characterization of the virus in rhesus macaque with the support of vet staff at UC Davis. X.C. prepared the figures with input from all authors. X.C. and V.G. wrote the manuscript with input from all authors. V.G. supervised all aspects of the work.

## ACKNOWLEDGMENTS

We thank members of the Gradinaru group for their assistance in this study: Yaping Lei for help with virus production, Miguel Chuapoco for discussion on the macaque experiment, Min Jee Jang and Cameron Jackson for attempt on FISH experiment, Elisha Mackey for mouse colony management, Zhe Qu for lab management, Patricia Anguiano for administrative assistance, and the entire Gradinaru group for discussions. We thank I. Antoshechkin and the Millard and Muriel Jacobs Genetics and Genomics Laboratory at Caltech for providing sequencing service. We thank the Beckman Institute Single-Cell Profiling and Engineering Center (SPEC) at Caltech and Sisi Chen for providing sequencing machines. We thank Annie W. Lam and Jost Vielmetter of the Beckman Institute Protein Expression Center at Caltech for Ly6a protein production and for providing surface plasmon resonance machines. We thank Sian Murphy at SWC’s Neurobiological Research Facility (NRF) for the injection of the rats. We thank Cassandra Tang-Wing and Chris Chamberlain at UCSD for the injection of the marmosets. We thank Michael Metke and Vikram Pal Singh at UCSD for the tissue collection from marmosets. We thank the staff at the California National Primate Research Center for the experiment in rhesus macaque.

We thank Catherine Oikonomou for help with manuscript editing. This work was primarily supported by grants from the National Institutes of Health (NIH) to V.G.: NIH Director’s New Innovator DP2NS087949 and PECASE, NIH BRAIN Armamentarium 1UF1MH128336-01, NIH Pioneer 5DP1NS111369-04 and SPARC 1OT2OD024899. Additional funding includes the Vallee Foundation (V.G.), the Moore Foundation (V.G.), the CZI Neurodegeneration Challenge Network (V.G. and C.E.), and the NSF NeuroNex Technology Hub grant 1707316 (V.G.), the Heritage Medical Research Institute (V.G.) and the Beckman Institute for CLARITY, Optogenetics and Vector Engineering Research (CLOVER) for technology development and dissemination (V.G.), NIH BRAIN UG3MH120095 (J.T.T, V.G.), CNPRC base grant (NIH P51 OD011107, A.F.), the Swiss National Science Foundation (310030_188952, A.K), the Synapsis (grant 2019-PI02, A.K.), the Swiss Multiple Sclerosis Society (A.K.). C.E. is an investigator of the Howard Hughes Medical Institute.

## MATERIALS AND CORRESPONDENCE

Correspondence to Viviana Gradinaru.

## METHODS

### A. Plasmids

#### 1. Library preparation

The plasmids used for AAV library preparation were described previously (Ravindra Kumar et al. 2020; Deverman et al. 2016) (plasmid would be deposited at Addgene). Briefly, plasmid rAAV-ΔCap-in-cis-Lox2 (Fig. 1A) was used for building the heptamer insertion (*7-mer-i*) AAV library. Plasmid pCRII-9Cap-XE was used as a PCR template for the DNA library generation. Plasmid AAV2/9-REP-AAP-ΔCap was used to supplement the AAV library during virus production.

#### 2. DNA construct for AAV characterization

The AAV capsid AAV-X1 was built by inserting 7-mer peptides between AAs 588-589 of AAV9 cap gene in the pUCmini-iCAP-PHP.B backbone (Ravindra Kumar et al. 2020). The AAV-PHP.V1 capsid was described previously (Ravindra Kumar et al, 2020, Addgene 127847). The AAV capsid AAV-X1.1 (Addgene 196836: will be available upon publication), AAV-X1.2, and AAV-X1.3 were built by substituting AAs 452-458 of AAV-X1. The AAV capsids AAV-X1.4, AAV-X1.5, and AAV-X1.6 were built by mutation of AA 272/386/503 in AAV-X1 to Alanine. The AAV capsid AAV1-X1 was built by inserting a 7-mer peptide between AAs 588-589 of the AAV1 cap gene in AAV1-Rep-Cap (Challis et al. 2019) (Addgene 112862).

For *in vivo* validation of AAV capsids, we packaged the vectors with a single-stranded (ss) rAAV genome: pAAV:CAG-eGFP, pAAV:CAG-tdTomato (a gift from Edward Boyden, Addgene plasmid # 59462). To make pAAV:CAG-eGFP-3xMir122-TS (Addgene ID: will be deposited on Addgene), 3 copies of the Mir122-TS were cut out from plasmid CAG-GCaMP6f-3x-miR204-5p-3x-miR122-TS (Challis et al. 2019). To make pAAV:CAG-Hevin-HA, Hevin-HA was synthesized as a gBlocks Gene Fragment (IDT) based off the sequence in the plasmid pAAV:GfaABC1D-Hevin, a gift from Cagla Eroglu’s Lab, and subcloned into the plasmid pAAV:CAG-eGFP by replacing the eGFP gene. To make pAAV:CAG-Ly6a, the Ly6a coding sequence from C57BL/6J was synthesized as a gBlocks Gene Fragment (IDT) and subcloned into the plasmid pAAV:CAG-eGFP by replacing the eGFP gene. pAAV-CAG-FXN-HA was chosen for the *ex vivo* slice study because it contains a ubiquitous CAG promoter and a HA-tagged endogenous human frataxin (FXN) protein and a unique 12bp barcode sequence. The barcode sequence was used to differentiate different capsid packaging the same construct during the next-generation sequencing (NGS) analysis.

#### 3. AAV capsid library generation

The round-1 (R1) and round-2 (R2) libraries were generated as described previously (Ravindra Kumar et al, 2020). Briefly, the R1 library involved a randomized 21-nucleotide (7xNNK mutagenesis) insertion between AAs 588-589 of the AAV9 capsid. The R2 library was built using a *synthetic pool* method (Ravindra Kumar et al. 2020). The R2 library was composed of an equimolar ratio of ∼4000 variants that were recovered from the tissues of interest in R1.

### B. Animals and subjects

All animal procedures in mice that were carried out in this study were approved by the California Institute of Technology Institutional Animal Care and Use Committee (IACUC), Caltech Office of Laboratory Animal Resources (OLAR), Cantonal Veterinary Office Zurich (license number ZH194/2020, 32869/2020), Duke Division of Laboratory Animal Resources (DLAR). All experimental procedures in rats were conducted at UCL according to the UK Animals Scientific Procedures Act (1986) and under personal and project licenses granted by the Home Office following appropriate ethics review.

All experimental procedures performed on marmosets were approved by the University of California, San Diego, Institutional Animal Care and Use Committee (IACUC) and in accordance with National Institutes of Health and the American Veterinary Medical Association guidelines. Two female animals and one male animal were used in this study and received intravenous injections of AAVs.

All experimental procedures performed on rhesus macaques were approved by the International Animal Care and Use Committee at the University of California, Davis and the California National Primate Research Center (CNPRC). One infant female animal was used in this study and received intravenous injections of AAVs.

All human neurosurgical tissue studies were approved by the Western Institutional Review Board. Human neurosurgical specimens were obtained with informed consent of patients that underwent neocortex resection for the treatment of temporal lobe epilepsy or for tumor removal. Specimens for research were not required for diagnostic purposes and were distal to the pathological focus of the surgical resections (Berg et al, 2021 Nature).

For all the experiments performed in this study, the animals were randomly assigned, and the experimenters were not blinded while performing the experiments unless mentioned otherwise.

#### C. *In vivo* selection and capsid library recovery

For capsid selection *in vivo*, the virus library was intravenously administered to male and female mice of various Cre transgenic lines (n=2-3 per Cre line) at 3×10^11^ vg per mouse in R1 selection, and at 3×10^11^ vg per mouse in R2 selection. Two weeks post injection, mice were euthanized, and the organs of interest were harvested and snap-frozen on dry ice. The tissues were stored at −80°C for long-term. To recover capsids from the tissue, the rAAV genome extractions from tissues were processed using Trizol, and the rAAV genomes were recovered by Cre-dependent PCR or Cre-independent PCR as previously described (Ravindra Kumar et al. 2020). The AAV DNA library, virus library and the libraries recovered from tissue post *in vivo* selection were processed for NGS as also described previously (Ravindra Kumar et al, 2020).

#### D. Characterization of AAV vectors across models

##### 1. AAV vector production

The AAV vectors were produced using an optimized vector production protocol (Challis et al, 2019). Plasmids were transfected into HEK 293T cells (ATCC) using polyethylenimine. We collected the medium 72 hours after the transfection and harvest both the cell and medium 120 hours after transfection. We precipitated the viral particles from the medium with 40% polyethylene in 500 mM NaCl and combined them with cell pellets for processing. The cell pellets were later suspended with 100 U/mL of salt-activated nuclease in 500 mM NaCl, 40 mM Tris, 2.5 mM MgCl2. AAVs were then harvested from the cell lysates with iodixanol step gradients and ultracentrifugation. Afterward, AAVs were concentrated with Amicon filters and formulated in sterile PBS solution. Transmission electron microscopy (TEM) was used to examine the purity of the prep. Virus titer was measured by qPCR. The average yield was ∼ 1×10^12^ vg per plate. BR1:CAG-eGFP was purchased from Signagen (SL116035).

##### 2. AAV vector administration in mice and tissue harvest

For the cell-type profiling of the novel AAVs in mice, the AAV vectors were injected intravenously via the retro-orbital route to 6-8 week old adult mice at a dose of 0.1-1×10^12^ vg per mouse. The retro-orbital injections were performed as described previously (Yardeni et al. 2011; Challis et al. 2019). The expression times were ∼3 weeks from the time of injection. The dosage and expression time were kept consistent across different experimental groups unless noted otherwise. To harvest the tissues of interest, the mice were anesthetized with Euthasol (pentobarbital sodium and phenytoin sodium solution, Virbac AH) and transcardially perfused using 30 – 50 mL of 0.1 M phosphate buffered saline (PBS) (pH 7.4), followed by 30 – 50 mL of 4% paraformaldehyde (PFA) in 0.1 M PBS. The organs were collected and post-fixed 24-48 h in 4% PFA at 4°C. Following this, the tissues were washed with 0.1 M PBS twice and stored in fresh PBS-azide (0.1 M PBS containing 0.05% sodium azide) at 4°C.

In the Hevin experiment, 4-month-old Hevin KO mice (Kucukdereli et al. 2011) were retro-orbitally injected with either AAV-X1.1:CAG-Hevin-HA or AAV-X1.1:CAG-eGFP (1E12 vg per mouse). After 3 weeks, the mice were perfused and brains were extracted for synapse assay. In the experiment with pericyte-deficient mice, 4-5 month-old PDGFB-retention motif knock out mice (*Pdgfb*^*ret/ret*^) in a C57BL6/J genetic background (Lindblom et al. 2003) were used. 3E11 vg per mice for X1 and 1E12 vg per mice for X1.1 were injected into mice via the tail vein. 3 weeks post-injection, the anaesthetized animals were perfused for 1–2 min with PBS, followed by 5 min perfusion with 4% PFA in PBS, pH 7.2. Brains were collected and post-fixed in 4% PFA in PBS, pH 7.2 at 4 °C for 6 h.

##### 3. AAV vector administration in rat and tissue harvest

Female rats were used (150-200g) for the experiments. 1E13 vg of the virus was delivered intravenously through the tail vein under light anesthesia. The injected volume was 0.5 ml containing the required number of particles. After 21 days the animals were perfused using 4% PFA solution and PBS. Brains were collected.

##### 4. AAV vector administration in marmoset and tissue harvest

Marmoset monkeys were anesthetized using an intramuscular Ketamine (20 mg/kg) and Acepromazine (0.5 mg/kg) injection. An intravenous catheter was placed in the saphenous vein of the hind leg and flushed with ∼2 mL of LRS (Lactated Ringer’s solution) for 2 min. Viruses were pooled together in a single syringe (∼500-900 µL) and infused at a rate of 200 µL/min into the catheter. Following the infusion, the catheter was flushed with ∼3 mL of LRS for 2 min and removed. The animal was then returned to a recovery cage.

Following an incubation period of 4-6 weeks post viral injection, the animals were euthanized by injecting pentobarbital intraperitoneally. Two researchers worked in parallel to harvest the tissue to limit degradation as quickly as possible. Each organ – brain, lungs, kidneys, etc - was removed and separated into two parts. One half of the tissue was flash-frozen in 2-methylbutane that was chilled with dry ice to preserve mRNA and DNA in the harvested tissues. The other half of the tissue was fixed in 4% PFA solution for estimation of protein expression. Flash-frozen tissue samples were transferred to a -80°C freezer, while PFA-fixed tissue samples were stored in a 4°C fridge.

##### 5. AAV vector administration in macaque and tissue harvest

One female rhesus macaque was injected within 10 days of birth. Prior to injection, the animal was anesthetized with ketamine (0.1 mL) and the skin over the saphenous vein was shaved and sanitized. AAVs (2.5×10^13^ vg/kg) were slowly infused into the saphenous vein for ∼1 min in < 0.75 mL of 0.1 M PBS. The animal was monitored while they recovered from anesthesia in their home enclosure, and daily for the remainder of the study. The monkey was individually housed within sight and sound of conspecifics.

Tissues were collected 4 weeks post AAV administration. The animal was deeply anesthetized and euthanized using sodium pentobarbital in accordance with guidelines for humane euthanasia of animals at the CNPRC. The whole body was perfused with ice cold RNase-free 0.1 M PBS. The brain was removed from the skull and blocked into 4 mm thick slabs in the coronal plane. Brain slabs and organs were subsequently post-fixed in 4% PFA for 48 h. One hemisphere of the animal was cryoprotected in 10%,15%, and 30% sucrose in 0.1 M PBS.

##### 6. AAV vector administration in *ex vivo* human and non-human primate brain slices

All procedures involving non-human primates conformed to the guidelines provided by the US National Institutes of Health. *Ex vivo* brain slice culture experiments were performed on temporal cortex tissue from adult *Macaca nemestrina* or *Macaca mulatta* animals housed at the Washington National Primate Research Center. We obtained these brain samples through the Tissue Distribution Program operating under approved University of Washington IACUC protocol number 4277-01. All human neurosurgical tissue studies were approved by the Western Institutional Review Board. Human neurosurgical specimens were obtained with informed consent of patients that underwent neocortex resection for the treatment of temporal lobe epilepsy or for tumor removal. Specimens for research were not required for diagnostic purposes and were distal to the pathological focus of the surgical resections (Berg et al. 2021).

Human and macaque brain slices were prepared using the NMDG protective recovery method (Ting et al., 2014; Ting et al., 2018). Human neurosurgical tissue or macaque brain tissue specimens were placed in carbogenated NMDG artificial cerebral spinal (ACSF) solution containing (in mM): 92 NMDG, 2.5 KCl, 1.25 NaH2PO4, 30 NaHCO3, 20 HEPES, 25 glucose, 2 thiourea, 5 Na-ascorbate, 3 Na-pyruvate, 0.5 CaCl2.4H2O and 10 MgSO4.7H2O. Brain slices were prepared on a VF-200 Compresstome at 300 μm thickness using a zirconium ceramic blade (EF-INZ10, Cadence) and then underwent warmed recovery in carbogenated NMDG aCSF at 32-34*C for 12 minutes. Human and macaque brain slices were placed on membrane inserts in 6 well sterile culture plates, and wells were filled with slice culture medium consisting of 8.4 g/L MEM Eagle medium, 20% heat-inactivated horse serum, 30 mM HEPES, 13 mM D-glucose, 15 mM NaHCO3, 1 mM ascorbic acid, 2 mM MgSO4.7H2O, 1 mM CaCl2.4H2O, 0.5 mM GlutaMAX-I, and 1 mg/L insulin (Ting et al., 2018). The slice culture medium was carefully adjusted to pH 7.2-7.3, osmolality of 300-310 mOsmoles/Kg by addition of pure H2O and then sterile-filtered. Culture plates were placed in a humidified 5% CO2 incubator at 35°C and the slice culture medium was replaced every 2-3 days until end point analysis. 1-3 hours after plating, brain slices were infected by direct application of ∼2.5×10^10^ vg of concentrated AAV viral particles distributed over the slice surface.

##### 7. AAV vector characterization in cell cultures

Human brain microvascular endothelial cells (HBMEC) (ScienCell Research Laboratories, cat. no. 1000) were cultured as per the instructions provided by the vendor. The viral vectors packaging ssAAV: CAG-eGFP were added to the cell culture at the multiplicity of infection (MOI) of either 5E4 or 5E3 per well (6 wells per dose per vector). The culture was assessed for fluorescence expression at 1-day post infection. In the *in vitro* Ly6a experiment, HEK293T cells (ATCC, CRL-3216) were cultured in 6-well plate, Ly6a was transiently expressed in HEK293T cells by transfecting each well with 2.53 µg plasmid DNA. Receptor-expressing cells were transferred to 96-well plates and transduced with AAV variants at the MOI of 3E4 or 3E3 per well. The culture was assessed for fluorescence expression at 1-day post infection. HeLa cells (ATCC, CCL-2), U87 cells (ATCC, HTB-14) and IMR32 cells (ATCC, CCL-127) were cultured as per the instructions provided by the vendor. The viral vectors packaging ssAAV: CAG-eGFP were added to the cell culture at the MOI of 5E4 per well (3 wells per dose per vector). The culture was assessed for fluorescence expression at 1-day post infection.

#### E. Immunohistochemistry and Imaging

In the AAV characterization experiment in mice, tissue sections, typically 100-µm thick, were first incubated in the blocking buffer (10% normal donkey serum (NDS), 0.1% Triton X-100, and 0.01% sodium azide in 0.1 M PBS, pH 7.4) with primary antibodies at appropriate dilutions (Anti-CD31 from Histonova Cat # DIA-310, diluted 1:100; Anti-podocalyxin from R&D Systems Cat # AF1556, diluted 1:100; Anti-CD13 from AbD Serotec, Cat # MCA2183EL, diluted 1:100; Anti-CNN-1 (Calponin 1) from Abcam, Cat # Ab46794, diluted 1:100; Anti-Collagen-IV, from AbD, Serotec Cat # 2150-1470, diluted 1:300; Anti-GFAP from Invitrogen, Cat # 13-030, diluted 1:600; Anti-HA from Roche, Cat# 11867423001, diluted 1:200; Anti-GFP from Aves Labs, Cat# GFP-1020, diluted 1:500; Anti-GLUT1 from Millipore, Cat# 07-1401, diluted: 1:500; Anti-S100β from Abcam, Cat# ab52642, diluted: 1:500; Anti-NeuN from Abcam, Cat# ab177487, diluted: 1:500) for 24 h at room temperature (RT) on a rocker. After primary antibody incubation, the tissues were washed 1-3 times with wash buffer 1 (0.1% Triton X-100 in 0.1 M PBS buffer, pH 7.4) over a period of 5 – 6 h in total. The tissues were then incubated in the blocking buffer with the secondary antibodies at appropriate dilutions for 12-24h at RT and then washed 3 times in 0.1 M PBS over a total duration of 5-6 h. When performing DNA staining, the tissues were incubated with 4’, 6-diamidino-2-phenylindole (DAPI) (Sigma Aldrich, 10236276001, 1:1000) in 0.1 M PBS for 15 min followed by a single wash for 10 min in 0.1 M PBS. The DAPI and/or antibody-stained tissue sections were mounted with ProLong Diamond Antifade Mountant (ThermoFisher Scientific, P36970) before imaging them under the microscope. The images were acquired with a Zeiss LSM 880 confocal microscope using the following objectives: Plan-Apochromat 10× 0.45 M27 (working distance 2.0 mm), and Plan-Apochromat 25× 0.8 Imm Corr DIC M27 multi-immersion. The liver images were acquired with a Keyence BZ-X700 microscope using a 10x objective. The images were then processed in the following image processing softwares: Zen Black 2.3 SP1 (for Zeiss confocal images) and BZ-X Analyzer (for Keyence images).

In the rat experiment, whole brains were imaged using serial section two-photon microscopy (Mayerich, Abbott, and McCormick 2008; Ragan et al. 2012). Our microscope was controlled by ScanImage Basic (Vidrio Technologies, USA) using BakingTray, a custom software wrapper for setting up the imaging parameters (Campbell 2020). 50-60um slices were cut and 7-9 optical planes were imaged using a 16x objective. Images were assembled using StitchIt (Campbell, Blot, and lguerard 2020).

In the Hevin experiment, for synaptic puncta analysis of mouse primary visual cortex (area V1), brains were cryosectioned at 25 µm using Leica CM3050S (Leica, Germany). Tissue sections were washed and permeabilized in TBS with 0.2% Triton-X 100 three times at room temperature followed by blocking in 5% Normal Goat Serum (NGS) for 1 hr at room temperature. To label pre and postsynaptic proteins, VGluT2 (Synaptic Systems; Cat# 135 404) and PSD95 (Thermo Fisher; Cat# 51-6900) antibodies were used, respectively. Primary antibodies were diluted in 5% NGS containing TBST and incubated overnight at 4 °C. Secondary antibodies (Alexa Fluor conjugated; Invitrogen) were added in TBST with 5% NGS for 2hr at room temperature. Slides were mounted in Vectashield with DAPI (Vector Laboratories, CA). Images were acquired with Olympus FV 3000 inverted confocal microscope using high magnification 60x objective plus 1.64x optical zoom z-stack images containing 15 optical sections spaced 0.33μm apart. During the post processing of captured images, each z-stack was converted into 5 maximum projection images (MPI) by combining three optical sections using ImageJ software. The number of co-localized excitatory thalamocortical (VGluT2/PSD95) synaptic puncta were obtained using the ImageJ plugin Puncta Analyzer (Ippolito and Eroglu 2010).

In the pericyte-deficient mice experiment, coronal vibratome sections (60 μm) of brains were cut using the Leica VT1000S. Free floating brain sections were incubated in the blocking/permeabilization solution (1% bovine serum albumin, 0.5% Triton X-100 in PBS) overnight at 4 °C, followed by incubation in primary antibody solution for two nights at 4 °C, and subsequently in secondary antibody solution, overnight at 4 °C. Sections were incubated with DAPI (4’,6-Diamidino-2-phenylindole dihydrochlorid) solution(D9542, Sigma-Aldrich, diluted 1:10000) for 7 minutes at RT and subsequently mounted in ProLong Gold Antifade mounting medium (cat. #P36930). Life Technologies). Images were taken with Leica SP8 inverse, 20 X objective PL APO CS (NA 0.7) (Leica Microsystems). Widefield images were generated using Slidescanner Zeiss Axio Scan.Z1 (Leica Microsystems). Image processing was done using Fuji and Zen2. All confocal images are represented as maximum intensity projections.

In the marmoset and macaque experiment, coronal sections (100 μm) of brains were cut using the Leica VT1000S. Sections (50-100 μm) of gut, DRG and spinal cord were cut using a cryostat (Leica Biosystems). Tissue were stained with relevant antibody following the similar protocol in the mouse characterization experiment. The images were acquired with a Zeiss LSM 880 confocal microscope using the following objectives: Plan-Apochromat 10× 0.45 M27 (working distance 2.0 mm), and Plan-Apochromat 25× 0.8 Imm Corr DIC M27 multi-immersion. The images were then processed in the Zen Black 2.3 SP1 (for Zeiss confocal images).

In the primate *ex vivo* slice culture experiments, following 7-10 days post virus infection, temporal cortex brain slices were fixed in 4% PFA for 24 hours at 4*C. After PFA fixation, the slices were transferred into 30% sucrose in water for >24 hours and then sub-sectioned on a sliding microtome to 15-30 μm for immunostaining using the following antibodies: mouse anti-HA (Biolegend catalog #901513, 1:1000) and rabbit anti-NeuN (Millipore catalog #ABN78, 1:2000). In a subset of IHC experiments we performed co-immunostaning with additional cell type marker antibodies including the following: rabbit anti-GLUT1 (Millipore catalog #07-1401, 1:1000), rabbit anti-Olig2 (Abcam catalog #AB9610, 1:1000), and mouse anti-S100β (Millipore catalog #S2532, 1:1000). We also switched from mouse anti-HA to rat anti-HA (Roche catalog #3F10, 1:1000) to circumvent antibody cross reactivity issues observed with Olig2 and other cell type markers. Secondary antibodies included Goat anti-mouse, Goat anti-rabbit, or Goat anti-rat Alexa Fluor conjugated antibodies (A488, A555, A647 from ThermoFisher) as needed. Slices were incubated in 1ug/mL DAPI solution and mounted on 1×3 inch slides with Prolong Gold mounting medium. Region of interest (ROI) in the neocortical grey matter spanning L3/4/5 were imaged on an Olympus FV3000 confocal microscope using 405nm, 561nm, and 640nm laser lines. Z-stack images were acquired at 1um step sized through the slice thickness and collapsed to made maximum intensity projection images. Every attempt was made to select comparable ROIs in the brain slices that best captured viral reporter expression patterns and to image at matched settings to directly compare across the capsid variants. In some cases, images needed to be pseudo-colored post-acquisition for best effect of illustrating signal overlap in two channel merged images (green and magenta with overlap in white).

In the RNAScope FISH and antibody labeling on ex vivo slice culture experiments, 350 µm thick macaque cortical slices were obtained, cultured and transduced as above. After eight days in culture, slices were fixed in 4% PFA (Electron Microscopy Sciences) at 4°C for 12 hours and cryoprotected in 30% sucrose. Thick slices were subsection 15 µm on a freezing sliding microtome (Leica SM2000R) and stored in 1 x phosphate buffered saline (PBS) + 0.1% sodium azide at 4°C. RNAscope Multiplex Fluorescent v2 (Advanced Cell Diagnostics) was performed, and if not otherwise noted, buffers or solutions were obtained from the ACD v2 kit. Sections were washed in 1 x PBS, mounted onto slides, and dried at 60°C. Samples were fixed in 4% paraformaldehyde (PFA) for 15 min at 4°C followed by a series of five minute washes in solutions with increasing ethanol concentrations (50%, 70%, 100%, 100%). Hydrogen peroxide was applied to the sections, incubated at RT for 10 minutes, and washed in 1x PBS twice for 1 min. Samples were incubated in Target Retrieval Reagent for 5-7min at 100°C. Slides were rinse in deionized water and incubated in 100% ethanol for 3 minutes and air-dried. Hydrophobic barrier around the tissue was created with an ImmEdge Pen (Vector Laboratories, H-4000). Samples were treated with Protease III for 30 minutes at 40°C and then washed twice in 1x PBS for 1 min at RT. We used a C1 probe designed by ACD against SYFP2 for detection of the EGFP reporter mRNA. Sections were incubated in this probe for 3 hours at 40°C. Sections were then washed twice in 1x Wash Buffer for 2 minutes each at RT, then incubated in 5x Sodium Chloride Sodium Citrate (SSC) (Invitrogen) overnight at RT. Sections were incubated in Amp1 reagent for 30 min at 40°C then washed twice in 1x Wash Buffer for 2 min at RT. Sections were then incubated in Amp2 reagent for 30 min at 40°C, then washed twice in 1x Wash Buffer for 2 min at RT. Sections were then incubated in Amp3 reagent for 15 min at 40°C, then washed twice in 1X Wash Buffer for 2 min at RT. Sections were then incubated in HRP-C1 at 40C for 15 min, washed twice in 1x Wash Buffer for 2 min at RT, then incubated in TSA Cy3 (Perkin Elmer, diluted 1:1500 in TSA Buffer) at for 30 min at 40C. Sections were then incubated in HRP Blocker at 40°C for 30 min and washed twice in 1x Wash Buffer for 2 min each at RT. Sections were washed in 1x PBS for 2 min at RT then fixed in 4% PFA for 15 min at RT. Sections were washed twice in 1x PBS for 5 min. Next, slides were incubated in a blocking buffer of 10% normal goat serum (Jackson ImmunoResearch) in PBS for 15 min at RT. Sections were incubated in primary antibody for 1 hour at RT: rabbit Anti-GLUT1 (Millipore), chicken Anti-GFP (Aves). Samples were then washed twice in 1x PBS for 5 min each, then incubated in secondary antibody for 1 hour at RT: goat anti-rabbit Alexa Fluor 488 (ThermoFisher), goat anti-chicken Alexa Fluor 488 (ThermoFisher). Samples were washed twice in 1x PBS for 5 min each, counterstained with DAPI for 30 seconds at RT, then mounted with Prolong Gold Antifade Mountant (ThermoFisher Scientific, P36930). Samples were imaged on an Olympus confocal FluoView3000 using a 30x silicone immersion objective or a Nikon Ti2 epifluorescence microscope with a 20x objective. Montage images were stitched on the Olympus software (FV31S-SW), stacks were Z-projected and analyzed using FIJI. Images from the Nikon are not montages, are from a single plane. All analysis was done in FIJI.

#### F. Protein production

Ly6a-Fc was produced in Expi293F suspension cells grown in Expi293 Expression Medium (Thermo Fisher Scientific) in a 37 °C, 5% CO2 incubator with 130 rpm shaking. Transfection was performed with Expifectamine according to manufacturer’s instructions (Thermo Fisher Scientific). Following harvesting of cell conditioned media, 1 M Tris, pH 8.0 was added to a final concentration of 20 mM. Ni-NTA Agarose (QIAGEN) was added to ∼5% conditioned media volume. 1 M sterile PBS, pH 7.2 (GIBCO) was added to ∼3X conditioned media volume. The mixture was stirred overnight at 4 °C. Ni-NTA agarose beads were collected in a Buchner funnel and washed with ∼300 mL protein wash buffer (30 mM HEPES, pH 7.2, 150 mM NaCl, 20 mM imidazole). Beads were transferred to an Econo-Pak Chromatography column (Bio-Rad) and protein was eluted in 15 mL of elution buffer (30 mM HEPES, pH 7.2, 150 mM NaCl, 200 mM imidazole). Proteins were concentrated using Amicon Ultracel 10K filters (Millipore) and absorbance at 280 nm was measured using a Nanodrop 2000 spectrophotometer (Thermo Fisher Scientific) to determine protein concentration.

#### G. Surface Plasmon Resonance (SPR)

Experiments were performed using a Sierra SPR-32 Pro (Bruker). Ly6a-Fc fusion protein in HBS-P+ buffer (GE Healthcare) was immobilized to a protein A sensor chip at a capture level of approximately 1200 -1500 response units (RUs). Two-fold dilutions of rAAV beginning at 4 × 10^12^ v.g. mL^-1^ were injected at a flow rate of 10 µl min^-1^ with a contact time of 240 s and a dissociation time of 600 s. The protein A sensor chip was regenerated with 10 mM glycine pH 1.5 after each cycle. Kinetic data were double reference subtracted.

#### H. Bulk sequencing for capsid enrichment in *ex vivo* tissue

*Ex vivo* NHP or human brain slice cultures were either infected with the pool of viruses with equivalent molar ratio or infected with PHP.eB packaging non CAG-FXN-HA construct. After 7 or 10 days *in vitro*, the tissues were snap frozen and shipped to Caltech on dry ice. Tissues were immediately stored at -80°C upon receipt. RNA was extracted from the whole tissue or half of the tissue following a slightly modified Phenol/Chloroform protocol (Goertsen et al. 2021). RNA was further cleaned by treating with DNase using RNA Clean & Concentrator-5 kit (Zymo), and reverse-transcription was done on up to 1 μg purified RNA using Superscript IV VILO Master Mix (ThermoFisher). If DNA was needed from the tissue, DNA was extracted from the other half of the tissue using QIAamp DNA Mini Kit (Qiagen) followed the manufacturer’s instructions.

The barcoded FXN region was recovered from the resulting cDNA library or DNA using primers of 5’-TGGACCTAAGCGTTATGACTGGAC-3’ and 5’-GGAGCAACATAGTTAAGAATACCAGTCAATC-3’ and PCR was performed using Q5 2x Master Mix (New England BioLabs) at 25 cycles of 98°C for 10s, 63°C for 15s, and 72°C for 20s. Each sample was run in up to 5 reactions using up to 50 ng of cDNA or DNA, each, as a template. After PCR, samples were purified using DNA Clean & Concentrator-25 kit (Zymo). The barcoded FXN region was further recovered and the adaptor sequence was added by performing PCR using primers of 5’-ACGCTCTTCCGATCTTGTTCCAGATTACGCTTGAG-3’ and 5’-TGTGCTCTTCCGATCTTGTAATCCAGAGGTTGATTATCG-3’ at 10 cycles of 98°C for 10s, 55°C for 15s, and 72°C for 20s. Samples were then purified using DNA Clean & Concentrator-25 kit. Index sets in the NEBNext Dual Index Primers (New England BioLabs) were carefully chosen and added to the barcoded FXN region by performing PCR at 10 cycles of 98°C for 10s, 60°C for 15s, and 72°C for 20s. To further separate the sequence for later next-generation sequencing (NGS), the PCR samples were run on a 2% low-melting-point agarose gel for separation and recovery of the 210bp band.

NGS was performed on an Illumina MiSeq Next Generation Sequencer (Illumina) using a 150-cycle MiSeq Reagent Kit v3 (Illumina) following the manufacturer’s procedure. All samples were pooled in equal ratio to a 4 nM library. 10% 20 pM PhiX control was spiked in to add diversity to the library. Demultiplexing was done by BaseSpace Sequence Hub and the barcode counting analysis was performed using in-house Python code. For each brain slice culture, the enrichment of capsid variants was calculated by the ratio of the counts of their corresponding barcode to the counts of the corresponding barcode to the internal control capsid (e.g. AAV9 or PHP.eB). To correct any potential error due to titer determination, PCR amplification, or sequencing, DNA from the same pool of virus that was used for the brain slice culture infection was extracted, amplified, and included in the MiSeq NGS and analysis. The enrichment of the capsid variants was then normalized by the input viral DNA in the pool.

#### I. QPCR and RT-PCR for vector genome and RNA transcript in *ex vivo* tissue

Quantitative PCR and quantitative RT-PCR to measure AAV variant transduction and expression in NHP slice culture. We cut physiological slices (300 microns thick) from the superior temporal gyrus from one adult (age 14 year and 1 month, weight 10.83 kg) M. mulatta animal housed at University of Washington National Primate facility. The animal had been planned for routine euthanasia and we collected the brain as part of the facility’s tissue distribution program. We sectioned a block of superior temporal gyrus and the slices were recovered as described by Ting et al. 2021, and then we cultured the slices on the air-liquid membrane interface as described in Mich et al. 2021. At 30 minutes following slice plating, we transduced the slices on their face with ∼1-2 uL. CAG-EGFP vectors, packaged in AAV9 or X1.1 (titers approximately 5e13 gc/mL). We performed transductions on three biological replicates for each vector. We refreshed the culture medium every 48 hours until tissue harvest at 8 days in vitro. At harvest, we imaged the slices to confirm transduction, and then bisected the slices, and quick-froze each slice half in a dry ice-ethanol bath, then stored them at -20 degrees C until nucleic acid processing.

One slice half (∼20 mg wet tissue weight) was processed to isolate DNA, and the other slice half was processed to isolate RNA for cDNA preparation. We isolated DNA using the Qiagen DNeasy Blood and Tissue Kit (Qiagen, catalog # 69504) with no modifications to the protocol, yielding approximately 3-5 ug total DNA per slice half. We isolated RNA using first TRIzol (Thermo Fisher Scientific, catalog #15596026) to isolate crude dilute RNA, which we then purified and concentrated using the PureLink RNA Mini Kit (Thermo Fisher Scientific, catalog # 12183018A). On the column of the PureLink RNA Mini Kit, we also performed DNA digestion by modifying the first wash as follows: we first washed with 350 uL of Wash Buffer 1, then added 80 uL of RNase-Free DNaseI in RDD buffer (Qiagen catalog # 79254) and incubated the column at room temperature for 15 minutes, then washed again with 350 uL of Wash Buffer 1, and proceeded with washing and elution as per the protocol. Typically, this technique yielded ∼1.5 ug DNA-free total RNA per slice half. We then performed first-strand cDNA synthesis from 400 ng total RNA in 20 uL reactions using Promega GoScript Reverse Transcription Kit (Promega, catalog # A5000).

For quantitative PCR we used 100 ng DNA in a 20 uL amplification reaction using the following Taqman probes (all from Thermo Fisher Scientific): EGFP-FAM probe (Assay ID Mr04097229_mr, catalog #4331182), and custom genomic reference probe CN2386-2-VIC (Assay ID ARH6DUK catalog #4448512, designed to target both M. mulatta and M. nemestrina). For quantitative RT-PCR we used 1 uL of cDNA (20 ng of original total RNA) in a 10 uL amplification reaction using the following probes: EGFP-FAM probe (same probe as used for DNA detection), and custom housekeeping reference probe GAPDH-VIC (Assay ID APAAHJR, catalog #4448508, designed to target both M. mulatta and M. nemestrina). We amplified and detected these reactions on a Roche Lightcycler II instrument alongside standard curves consisting of serial dilutions of plasmid DNA or purified PCR amplicons in the case of GAPDH. Then we extracted Cp values using the second derivative method, calculated the relative abundances of vector DNA/RNA with respect to internal reference probes, and normalized these ratios to those observed with AAV9 vector. In some cases, the EGFP-FAM probe was so abundant that it interfered with detection of the VIC reference probes; in these cases, the reactions were run in separate wells.

#### J. Data analysis

##### 1. Quantification of AAV transduction *in vivo*

The quantification of AAV transduction across tissues was carried out by manually counting fluorescent expression resulting from the AAV genome. ImageJ was used for this purpose.

##### 2. NGS data alignment, processing and analysis

The NGS data analysis was carried out using a custom data-processing pipeline with scripts written in Python (Ravindra Kumar et al. 2020) (https://github.com/GradinaruLab/mCREATE) and using plotting software such as Plotly, Seaborn, and GraphPad PRISM 7.05. The AAV9 capsid structure model was produced with PyMOL.

##### 3. Enrichment score calculation

The enrichment score for a variant was determined using the following formula:

Enrichment score of variant “x” in tissue1= log10 [(Enrichment of variant ‘X’ in the tissue1) / (Enrichment of variant ‘X’ in the virus pool)]

Enrichment of variant ‘X’ in the tissue1 = (Variant “x” RC in tissue library) / (Sum of variants N RC in tissue1)

Enrichment of variant ‘X’ in the virus pool = (Variant “x” RC in virus pool) / (Sum of variants N RC in virus pool)

Where N is the total number of variants in a library.

##### 4. Fold change calculation

The fold-change of a variant “x” to AAV9 = (The enrichment score of “x”) / (The enrichment score of AAV9).

---

**Supplementary Figure 1:**
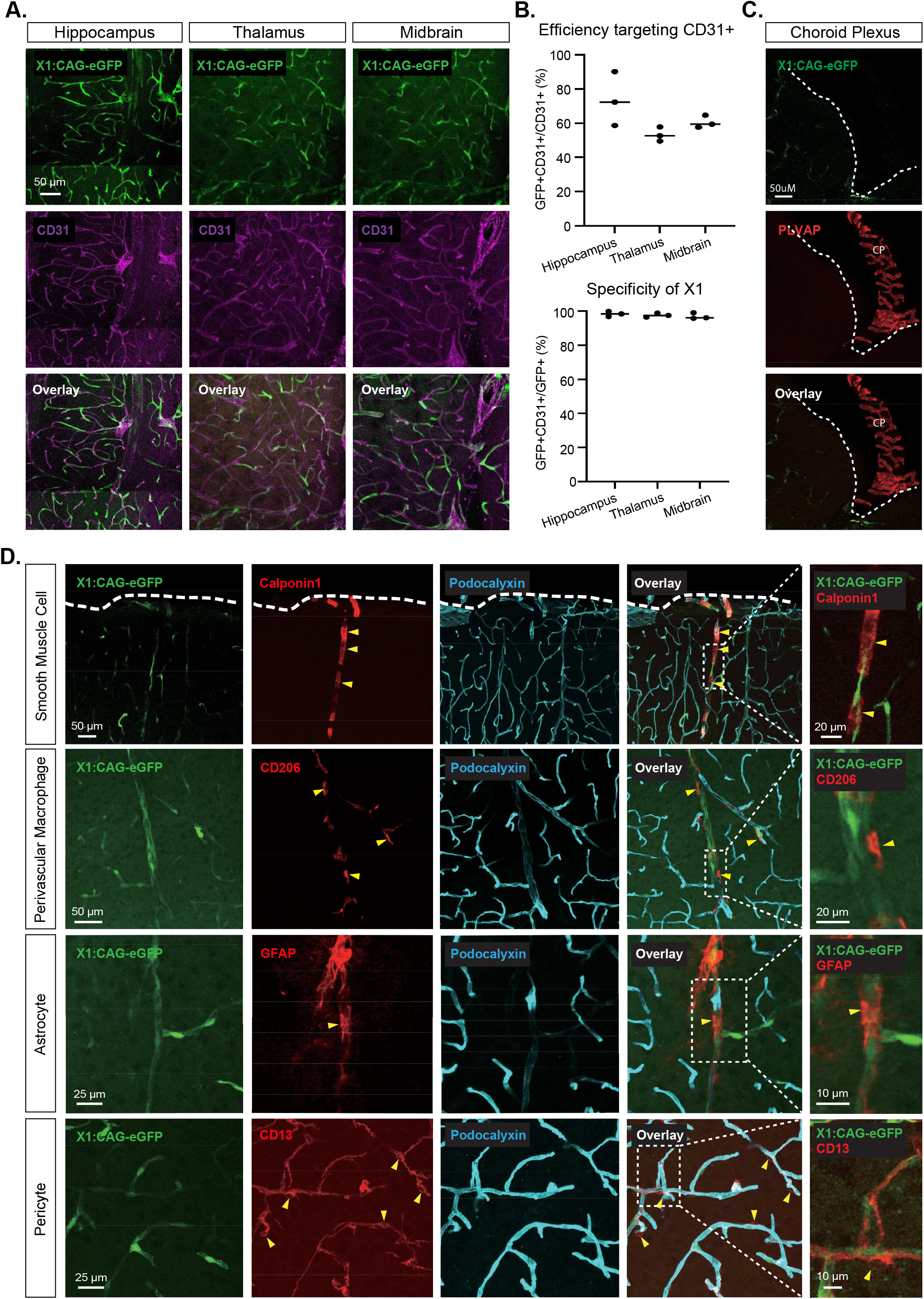
Detailed characterization of AAV9-X1 in the mouse brain. **A**. Representative images of AAV-X1 vector-mediated expression of eGFP (green) in the brain. The tissues were co-stained with GLUT1 marker (magenta) (scale bar: 50 µm). **B**. (Top) Percentage of AAV-mediated eGFP-expressing cells that overlap with GLUT1+ markers across brain regions, representing the efficiency of the vectors in targeting CD31+ cells. Each data point shows the mean ± s.e.m of 3 slices per mouse. (Bottom) Percentage of CD31+ markers in AAV-mediated eGFP-expressing cells across brain regions, representing the specificity of the vectors in targeting GLUT1+ cells. (n≥4 per group, ∼8 weeks old C57BL/6J males, 3×10^11^ vg IV dose per mouse, 3 weeks of expression). **C-D**. EGFP expression is seen only in endothelial cells possessing BBB characteristics, and not in other vascular cells. **C**. PLVAP (red)-positive endothelial cells in choroid plexus (CP) do not express eGFP. The ventricular border is indicated by the dashed line. **D**. Representative images of brain sections co-stained with endothelial cell marker (podocalyxin, in cyan) and (in red, with yellow arrowheads) markers for smooth muscle cells (calponin 1), perivascular macrophages (CD206), astrocytes (GFAP), and pericytes (CD13). The dashed lines indicate the cortical surface.

**Supplementary Figure 2:**
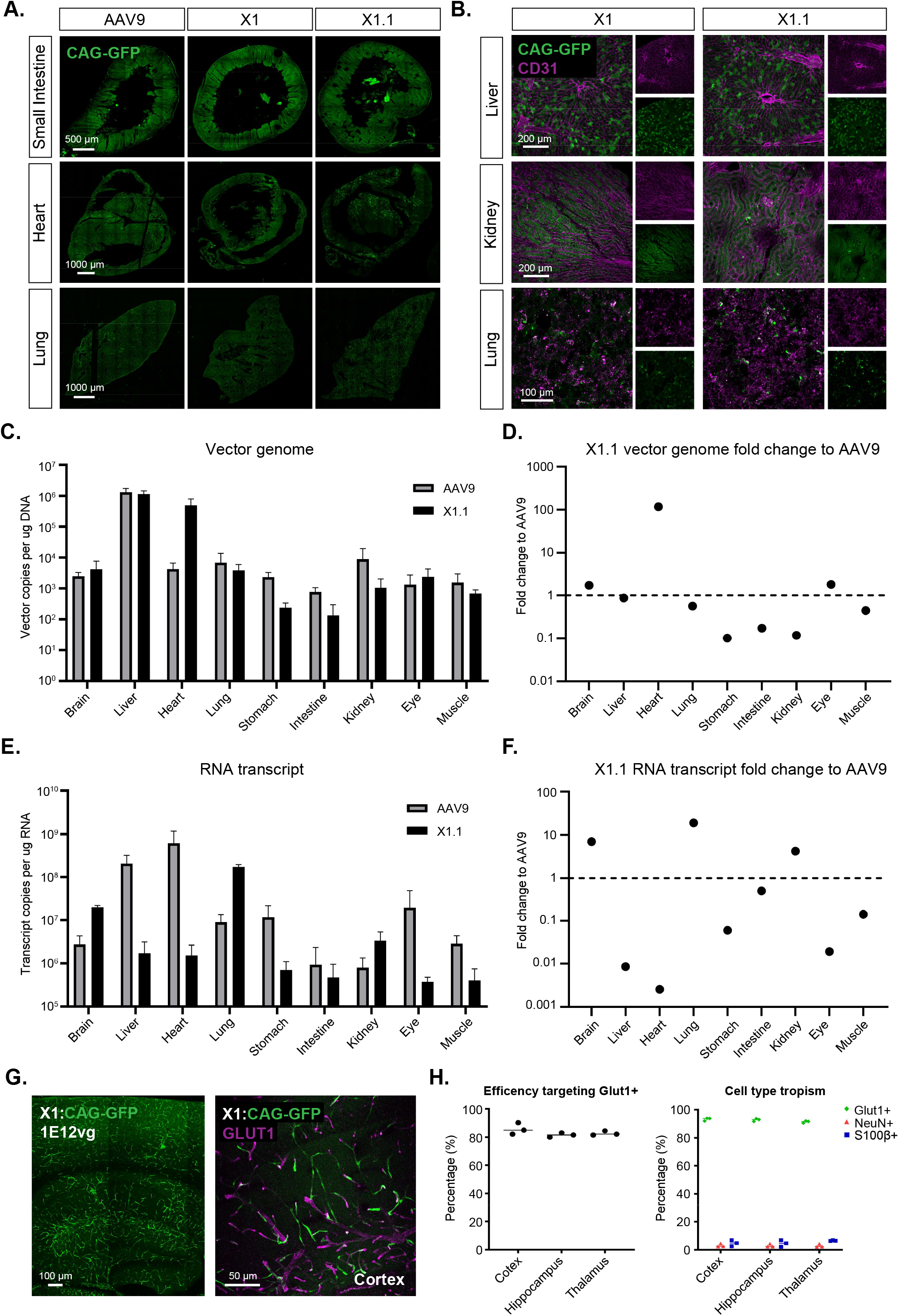
Biodistribution of AAVs across mice organs. **A**. Representative images of AAV9, AAV-X1, and AAV-X1.1 vector-mediated expression of eGFP in the small intestine (scale bar: 500 µm), heart (scale bar: 1000 µm) and lung (scale bar: 1000 µm) (n≥4 per group, ∼8 week-old C57BL/6J males, 3E11 vg IV dose per mouse, 3 weeks of expression). **B**. Representative images of AAV-X1, and AAV-X1.1 vector-mediated expression of eGFP in the liver (scale bar: 500 µm), kidney (scale bar: 1000 µm) and lung (scale bar: 1000 µm) (n≥4 per group, ∼8 week-old C57BL/6J males, 3E11 vg IV dose per mouse, 3 weeks of expression). The tissues were co-stained with CD31 (magenta). C. Vector genome of AAV9 or AAV-X1 per ug DNA across organs in mice following I.V. delivery (n=3 per group, ∼8 week-old C57BL/6J males, 3E11 vg IV dose per mouse, 3 weeks of expression). D. Vector genome of AAV-X1 fold change of vector genome of AAV9 across organs. (n=3 per group, ∼8 week-old C57BL/6J males, 3E11 vg IV dose per mouse, 3 weeks of expression). E. RNA transcript of AAV9 or AAV-X1 per ug RNA across organs in mice following I.V. delivery, GADPH was used to normalize across organs (n=3 per group, ∼8 week-old C57BL/6J males, 3E11 vg IV dose per mouse, 3 weeks of expression). F. RNA transcript of AAV-X1 fold change of RNA transcript of AAV9 across organs. (n=3 per group, ∼8 week-old C57BL/6J males, 3E11 vg IV dose per mouse, 3 weeks of expression).

**Supplementary Figure 3:**
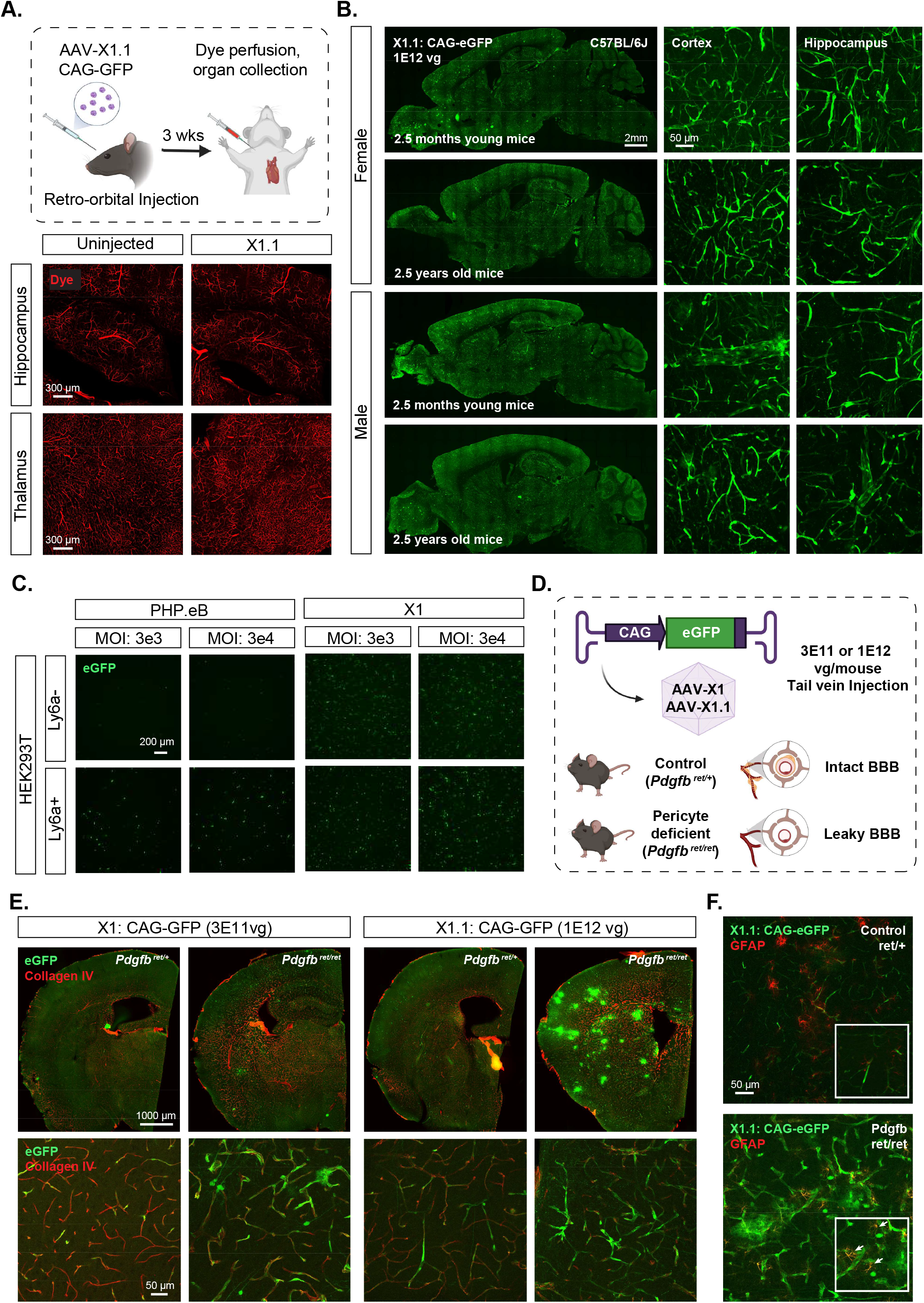
Engineered AAVs are independent of Ly6a and show different expression patterns in pericyte-deficient mice. **A**. (Top) Illustration of dye perfusion to evaluate the intactness of the BBB in AAV-injected mice. (Bottom) Representative images of dye staining (red) in hippocampus and thalamus (scale bars: 300 μm) (n=3 per group, 1E12 vg IV dose per mouse, 3 weeks of expression). **B**. Representative image of AAV-X1 vector-mediated expression of eGFP in the brains of both sexes and both young 2.5-month-old and aged 2.5-year-old C57BL/6J males (scale bars: 2 mm in whole brain and 50 μm in cortex/hippocampus) (n=3 per group, 1E12 vg IV dose per mouse, 3 weeks of expression). **C**. Representative images of AAV transduction in HEK293T cells with or without previous transfection of plasmids encoding Ly6a. (Packaged with ssAAV:CAG-eGFP, n= 3 per condition, 2-day expression, high dose: MOI 25000, low dose: MOI 2500). Scale bar: 200 µm. **D**. Illustration of AAV vector delivery to control mice and pericyte-deficient mice (Pdgfb ret/ret) for studying their transduction profile in BBB in different conditions (3E11 vg/mouse for X1, 1E12 vg/mouse for X1.1, tail vein injection, 3 weeks’ expression). **E**. Representative images of AAV-mediated expression of eGFP (green) in coronal sections of mouse brain (scale bar: 1000 µm), and zoomed-in images of tissue co-stained with collagen IV marker (red) (scale bar: 2 mm). **F**. Representative images of tissue co-stained with GFAP marker (red) (scale bar: 50 µm). Boxes show further zoomed-in views of astrocytes that have endfeet on the vasculature; white arrows highlight the colocalization of eGFP expression and GFAP marker in Pdgfb ret/ret mice.

**Supplementary Figure 4:**
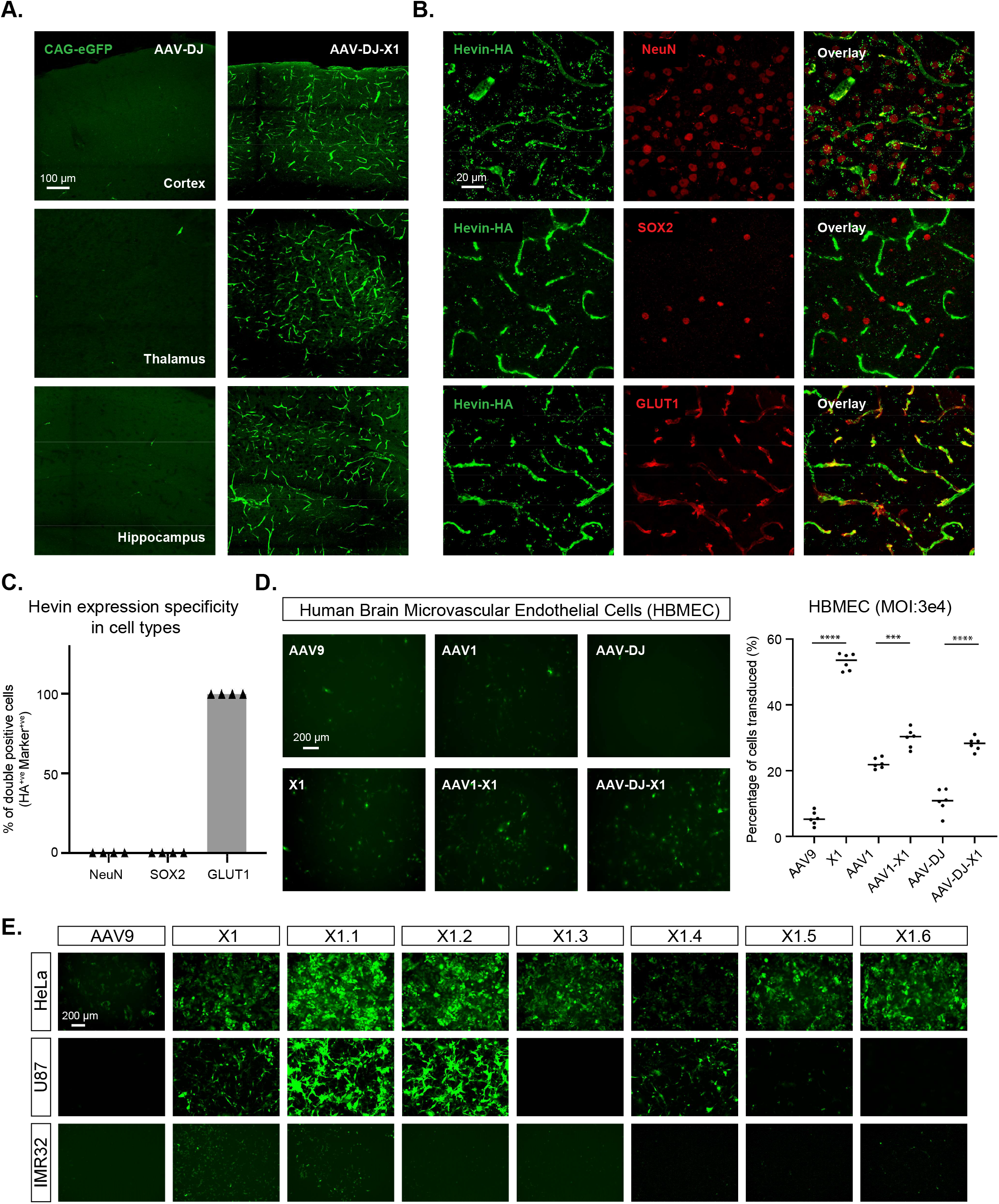
Hevin specificity amd engineered AAVs efficiently transduce human cells. **A**. Representative images of AAV-DJ and AAV-DJ-X1-mediated eGFP expression in the cortex, thalamus and hippocampus (scale bar: 100 µm) (C57BL/6J, n=3 per group, 3E11 vg IV dose per mouse, 3 weeks of expression). **B**. Representative images of AAV-X1.1 vector-mediated expression of Hevin in the brain. The tissues were co-stained with HA (green) and NeuN (red) or SOX2 (red) or GLUT1 (red) (scale bar: 30 µm). **C**. Quantification of AAV-X1.1-mediated Hevin expression across cell types. **D**. (Left) Representative images of AAV (AAV9, X1, AAV1, AAV1-X1, AAV-DJ, AAV-DJ-X1)-mediated eGFP expression (green) in Human Brain Microvascular Endothelial Cells (HBMECs). (AAVs packaged with ssAAV:CAG-eGFP, n= 6 per condition, 1 day expression). (Right) Percentage of cells transduced by the AAVs, one-way analysis of variance (ANOVA) non-parametric Kruskal-Wallis test (approximate P<0.0001), and follow-up multiple comparisons with uncorrected Dunn’s test are reported (P<0.0001 for AAV9 versus X1, P=0.0002 for AAV1 versus AAV1-X1, P<0.0001 for AAV-DJ versus AAV-DJ-X1; n=6 per group, each data point is the mean of 3 technical replicates, mean ± s.e.m is plotted). **E**. Representative images of AAV (AAV9, X1, X1.1, X1.2, X1.3, X1.4, X1.5, X1.6)-mediated eGFP expression (green) in HeLa cells, U87 cells, and IMR32 cells (AAVs packaged with ssAAV:CAG-eGFP, n= 3 per condition, 1 day expression, MOI: 50000).

**Supplementary Figure 5:**
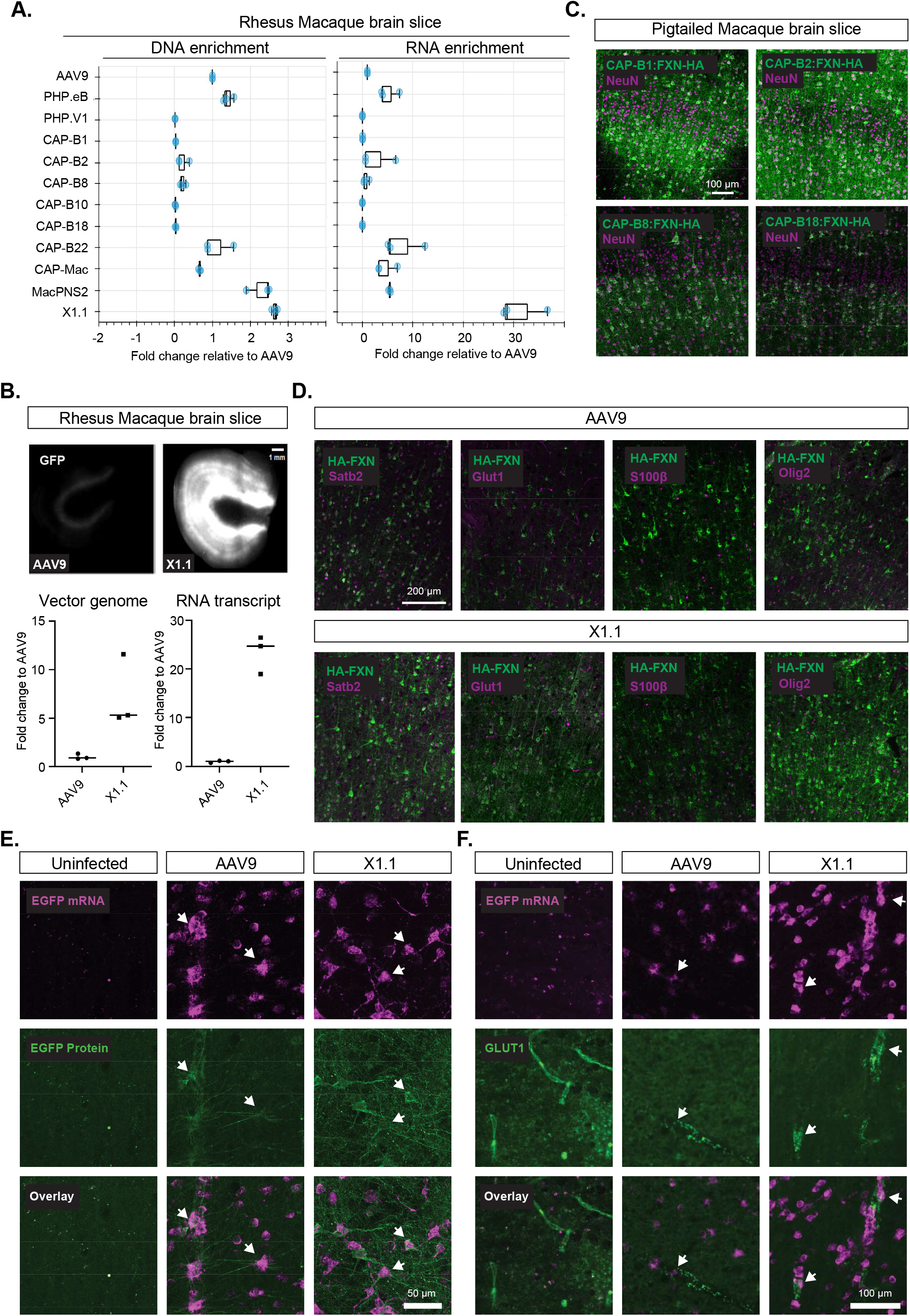
Engineered AAVs efficiently transduce human cell lines and *ex vivo* macaque slices. A. DNA and RNA level in rhesus macaque brain slices for AAVs, with DNA and RNA levels normalized to AAV9. **B**. (top) Representative images of AAV9 and X1.1-mediated CAG-GFP expression in *ex vivo* rhesus macaque brain slices. (bottom) Vector genome and RNA transcript of the AAVs in the slices, fold change to AAV9 was shown. **C**. Representative images of AAV (CAP-B1, CAP-B2, CAP-B8, CAP-B18)-mediated CAG-FXN-HA expression in *ex vivo* southern pig-tailed macaque brain slices. The tissues were co-stained with antibodies against HA (green) and NeuN (magenta). **D**. Representative images of AAV9 and X1.1-mediated CAG-FXN-HA expression in *ex vivo* southern pig-tailed macaque brain slices. The tissues were co-stained with antibodies against HA (green) and Satb2, GLUT1, S100β or Olig2 (magenta). **E, F**. mFISH analysis of EGFP expression in transduced cultured slices. Organotypic cortical slices were cultured from macaque brain, and were uninfected or infected by X1.1 or AAV9 encapsidated EGFP reporter vector. After eight days in culture, EGFP mRNA was resolved against EGFP protein (E) or GLUT-1 protein (F) in cortical white matter regions. Minimal background EGFP mRNA or protein was detected in the absence of virus (uninfected), EGFP mRNA positive cells often express EGFP protein (arrows, E), and EGFP mRNA positive cells are often found lining GLUT-1+ blood vessels (putative endothelial cells, or pericytes, arrows, F). Images are from stitched montages.

**Supplementary Figure 6:**
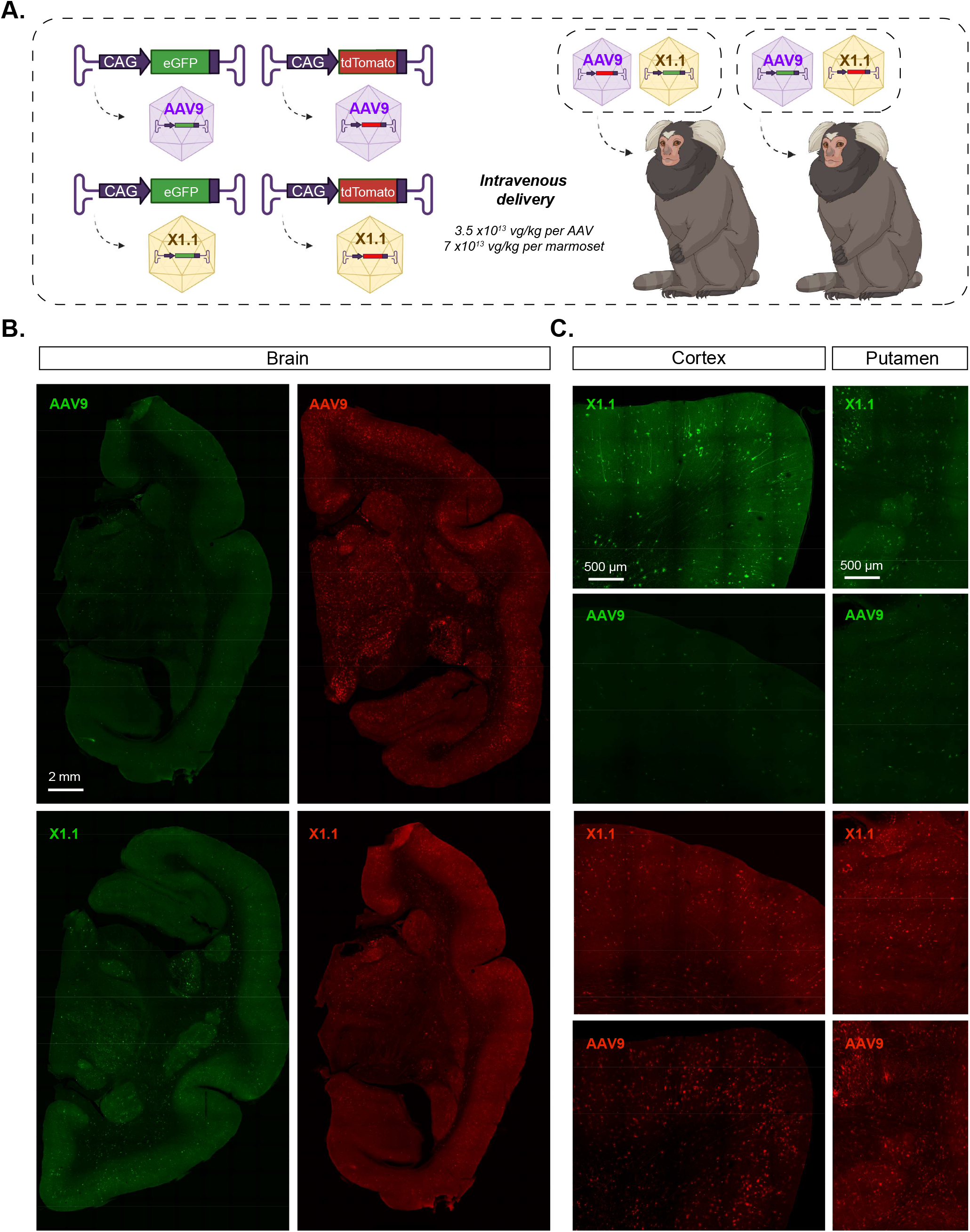
Engineered AAV transduces the central nervous system in marmoset similarly to AAV9. **A**. Illustration of AAV vector delivery to adult marmoset to study transduction across the CNS after 3 weeks of expression. The capsids (AAV9/X1.1) and their corresponding genomes (ssAAV:CAG-eGFP/tdTomato) are shown on the left. Two AAV vectors packaged with different colored fluorescent reporters were mixed and intravenously delivered at a total dose of 7×10^13^ vg/kg per adult marmoset (16 month-old *Callithrix jacchus*, i.e. 3.5×10^13^ vg/kg per AAV). Representative images of **B**., coronal brain sections of the midbrain (scale bar: 2 mm), and **C**., select brain areas: cortex and putamen (scale bar: 500 µm), showing AAV9 vector-mediated expression of eGFP (green) or tdTomato (red), X1.1-mediated expression of eGFP (green) and X1.1-mediated expression of tdTomato (red).

**Supplementary Figure 7:**
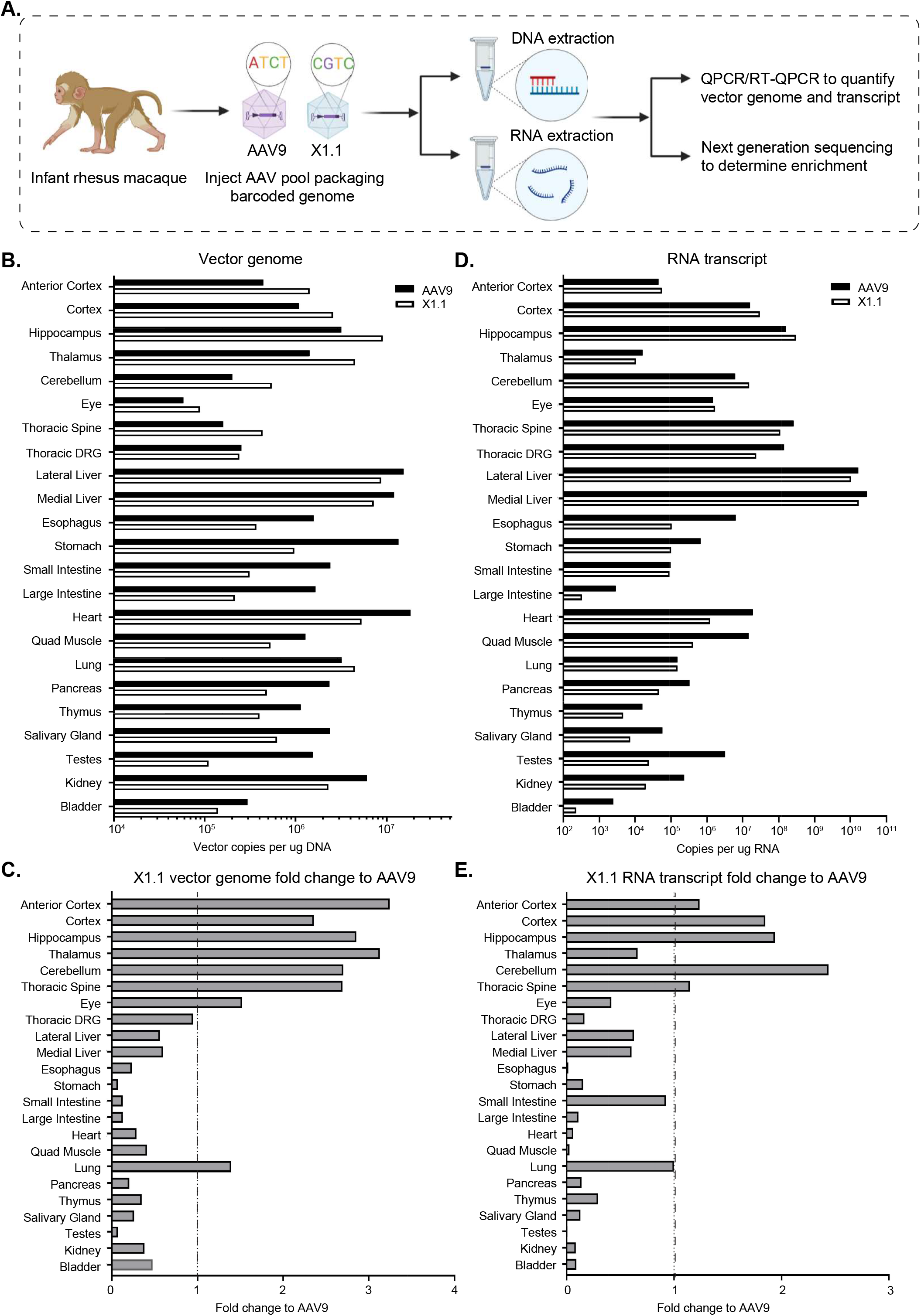
Biodistribution of engineered AAV in rhesus macaque following systemic delivery. **A**. Illustration of AAV vector delivery to rhesus macaque to study biodistribution after 4 weeks of expression. The capsids (AAV9/X1.1) packaged with CAG-GFP genome containing a unique barcode were mixed and intravenously injected at a dose of 2×10^13^ vg/kg per macaque (*Macaca mulatta*, injected within 10 days of birth, male, i.e. 2×10^13^ vg/kg per AAV). After 4 weeks of expression, tissues were collected and DNA/RNA extractions were performed. Then QPCR/RT-QPCR were performed to quantify vector genome and transcript. Next generation sequencing was performed to determine enrichment. **B**. Vector genome of AAV9 or AAV-X1 per ug DNA across organs in rhesus macaque following IV delivery. **C**. Vector genome of AAV-X1 fold change of vector genome of AAV9 across organs in rhesus macaque. **C**. RNA transcript of AAV9 or AAV-X1 per ug RNA across organs in rhesus macaque following I.V. delivery, GADPH was used to normalize across organs. F. RNA transcript of AAV-X1 fold change of RNA transcript of AAV9 across organs. (n=3 per group, ∼8 week-old C57BL/6J males, 3E11 vg IV dose per mouse, 3 weeks of expression).

**Supplementary Table 1:**
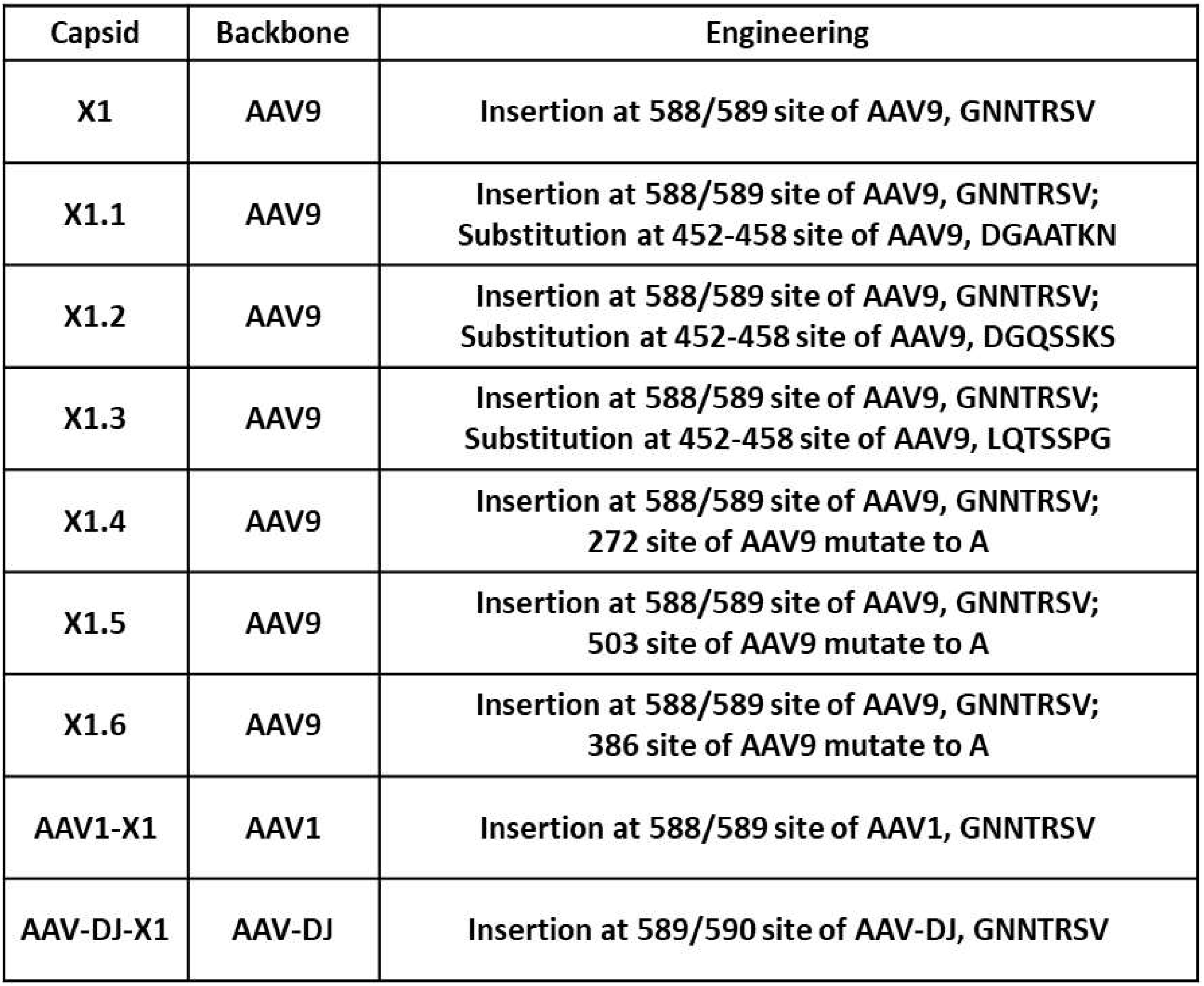
Summary of engineering approaches for the novel AAVs.

**Supplementary Table 2:** Summary of novel AAVs’ performance across models tested.

| Capsid | Model tested | Sex and age | Administration | Tropism |
| --- | --- | --- | --- | --- |
| X1 | Wildtype mice (C57BL/6J, BALB/CJ, FVB/NJ, CBA/J) | Both male and female, 6-8 weeks | Intravenously | Brain endothelial cell specific expression |
|  | Transgenic mice (PDGFB-retention motif knock out mice) | Both male and female, 6-8 weeks | Intravenously | Brain endothelial cell specific expression in PDGFB ret/+. Endothelial, neuron, astrocyte expression in PDGFP ret/ret |
|  | Human Brain Microvascular Endothelial cells | - | Direct application | Stronger expression comparing to controls including AAV9, AAV1, AAV2 etc. |
| X1.1 | Wildtype mice (C57BL/6J) | Both male and female, 6-8 weeks or 2 years old | Intravenously | Brain endothelial cell specific expression |
|  | Transgenic mice (Hevin-KO) | Male, 6-8 weeks | Intravenously | Brain endothelial cell specific expression |
|  | Transgenic mice (PDGFB-retention motif knock out mice) | Both male and female, 6-8 weeks | Intravenously | Brain endothelial cell specific expression in PDGFB ret/+, endothelial, neuron, astrocyte expression in PDGFP ret/ret |
|  | Rat | Both male and female, 6-8 weeks | Intravenously | Brain endothelial cell specific expression |
|  | Marmoset | Both male and female, 14 months | Intravenously | Neuron, astrocyte and endothelial cell expression |
|  | Rhesus Macaque | Both male and female, infant | Intravenously | Neuron and endothelial cell expression |
|  | Macaque Brain Slice | Both male and female, adult 2~4 years, | Direct application | predominantly neuron expression with lesser astrocyte, oligodendrocyte, and endothelial cell expression |
|  | Human Brain Microvascular Endothelial cells | - | Direct application | Stronger expression comparing to controls including AAV9, AAV1, AAV2 etc. |
|  | Human Brain Slice |  | Direct application | Stronger expression comparing to controls including AAV9 |

**Supplementary Video 1: Macaque hindbrain injected with AAV9 and AAV9-X1.1**. AAV9 packaging ssAAV:CAG-eGFP and AAV9-X1.1 packaging ssAAV:CAG-tdTomato were mixed and intravenously injected at a dose of 5×1013 vg/kg per macaque (Macaca mulatta, injected within 10 days of birth, female, i.e. 2.5×1013 vg/kg per AAV). Representative overview of macaque hindbrain.

